# Surface Binding Sites Orchestrate Long-Range Control of Active-Site Dynamics and Glucose Tolerance in β-glucosidase, BglB

**DOI:** 10.64898/2026.07.24.740578

**Authors:** Sneha Sahu, Sauratej Sengupta, Supratim Datta, Neelanjana Sengupta

**Author notes:** Corresponding author: Neelanjana Sengupta, Department of Biological Sciences, Indian Institute of Science Education and Research Kolkata Mohanpur 741246, West Bengal, India.;, Supratim Datta, Department of Biological Sciences, Indian Institute of Science Education and Research Kolkata Mohanpur 741246, West Bengal, India.

## Abstract

β-Glucosidases are essential for lignocellulosic biomass conversion, yet their industrial utility is limited by inhibition from glucose, the reaction product. Although glucose inhibition has been extensively documented, its molecular basis remains poorly understood. Here, we demonstrate that glucose inhibition in the GH1 β-glucosidase BglB from *Paenibacillus polymyxa* is mediated by multiple surface-exposed secondary binding sites (SBSs) that function as regulatory elements rather than passive glucose-binding patches. Kinetic analyses revealed a mixed mode of inhibition, with modest activation at low glucose concentrations followed by progressive inhibition at higher concentrations. Glucose association occurred through multiple SBSs in a concentration-dependent manner and induced localized rigidification, particularly at gatekeeper regions surrounding the active-site entrance. These surface interactions propagated through long-range coupling pathways, resulting in structural, energetic, and residue interaction network reorganization that reshaped the catalytic pocket. Functional interrogation of representative SBSs revealed distinct roles for individual sites. Mutation of a gatekeeper-associated SBS reduced local glucose association, increased glucose tolerance by more than 30%, and improved substrate affinity by approximately 38%, whereas disruption of a distal SBS compromised structural integrity and soluble protein production. Together, these findings establish a direct link between surface glucose recognition and active-site regulation, providing, to our knowledge, the first integrated evidence for the existence and functional significance of secondary glucose-binding sites in β-glucosidases. More broadly, this work identifies surface SBSs as promising targets for engineering glucose-tolerant and catalytically robust enzymes for biomass conversion and related biotechnological applications.

## Introduction

Lignocellulosic biomass is an abundant renewable resource, with cellulose, its primary constituent, produced at an estimated 1.5 trillion tonnes per year, making it a promising feedstock for sustainable biofuels. Efficient cellulose saccharification requires a synergistic cellulase cocktail made up of endoglucanases (EG) that cleave internal bonds, cellobiohydrolases (CBH) that release cellobiose from chain ends, and β-glucosidases (BG) that hydrolyze cellobiose into two glucose molecules^1,2^. However, many natural cellulase producers, such as *Trichoderma reesei*, express low levels of BG, which limits overall hydrolysis efficiency^3^. Additionally, the accumulation of glucose, a reaction product, further inhibits BG activity, creating a significant bottleneck in biomass hydrolysis by causing cellobiose buildup, which in turn inhibits CBH and EG activity.

To overcome these limitations, researchers have pursued protein engineering, enzyme supplementation, and computation-guided design to enhance glucose tolerance in BG. Computational analyses suggest that mutations that increase the mobility of flexible loops around the active site can hinder glucose entry and promote its withdrawal^4^. Combined experimental and computational studies on *Agrobacterium tumefaciens* 5A mutants indicate that conformational changes and additional glucose-binding sites correlate with improved tolerance^5^ and gatekeeper residues at tunnel entrances can control product release from the tunnel. Site-directed mutagenesis studies (for example, H184F in Bgl1B) and investigations of archaeal BGs support these themes^6, 7^. Recent literature underscores the complexity and variability of glucose-mediated inhibition: low glucose levels can sometimes stimulate BG activity while higher concentrations typically inhibit GH1 enzymes and reported inhibition strengths span millimolar to molar K_i_ values^8^. Contributing factors include glucose binding at secondary binding sites, conformational rearrangements, and differences between BG families (GH1 vs GH3)^7, 9–12^. *In silico* phase space sampling and molecular dynamics (MD) simulations offer atomic-level insights into transient, low-affinity interactions and ligand-induced conformational changes that are challenging to capture experimentally^13^. MD and related computational protocols can predict binding modes and affinities, identify residues governing stability and specificity, and prioritize mutations for experimental testing. Integrating simulation with experimental screening accelerates optimization of enzyme affinity, specificity, and thermostability – properties critical for robust biocatalysts^14–17^.

Here, we investigate the molecular basis of glucose-mediated inhibition in the β-glucosidase (EC 3.2.1.21) BglB from *Paenibacillus polymyxa* by integrating molecular simulations with biochemical and mutational analyses. We hypothesized that, beyond transient occupancy of the catalytic pocket, glucose associates with multiple surface-exposed secondary binding sites (SBSs) that collectively modulate enzyme structure and function. To test this hypothesis, we performed molecular dynamics simulations in glucose–water mixtures spanning a range of glucose concentrations and characterized glucose occupancy, protein dynamics, residue interaction networks, and thermodynamic responses. Guided by these computational insights, we experimentally evaluated representative SBSs to determine their contributions to glucose tolerance and catalytic function. Our results demonstrate that glucose association at surface SBSs is propagated through long-range structural communication pathways that remodel the active-site environment. These findings establish secondary binding sites as key determinants of product inhibition and identify them as promising targets for engineering glucose-tolerant β-glucosidases.

## Results and Discussion

### Effect of Glucose on BglB Activity

Since glucose is known to inhibit β-glucosidases, its effect on BglB kinetics was quantified using *p*-Nitrophenyl-β-D-glucopyranoside (*p*NPGlc) as the substrate under increasing concentrations of exogenous glucose. Steady-state kinetics were measured at 0, 0.05, 0.10, 0.15, 0.20, 0.30, and 0.50 M glucose. A slight increase in V_max_ was observed from 0.05 to 0.20 M, followed by a decline at higher concentrations. This change in V_max_ was accompanied by a continuous increase in K_m_ with increasing glucose concentration (Figure 1). Consistent with fold change in K_m_, catalytic efficiency (k_cat_/K_m_) decreased overall with increasing glucose concentration, reflected in reduced enzymatic activity. Enzymatic activity showed a transient enhancement at low glucose (+7% at 0.05 M), followed by a reduction beyond ∼0.15-0.20 M, decreasing by ∼20% at 0.1 M and further decreased to 27% at 1 M (Figure S1). Global fitting yielded an inhibition constant K_i_ of 0.09 M and an α value of 58.19 (>1), consistent with a mixed inhibition model in which glucose interacts with both the free enzyme and the enzyme-substrate complex.

**Figure 1.**
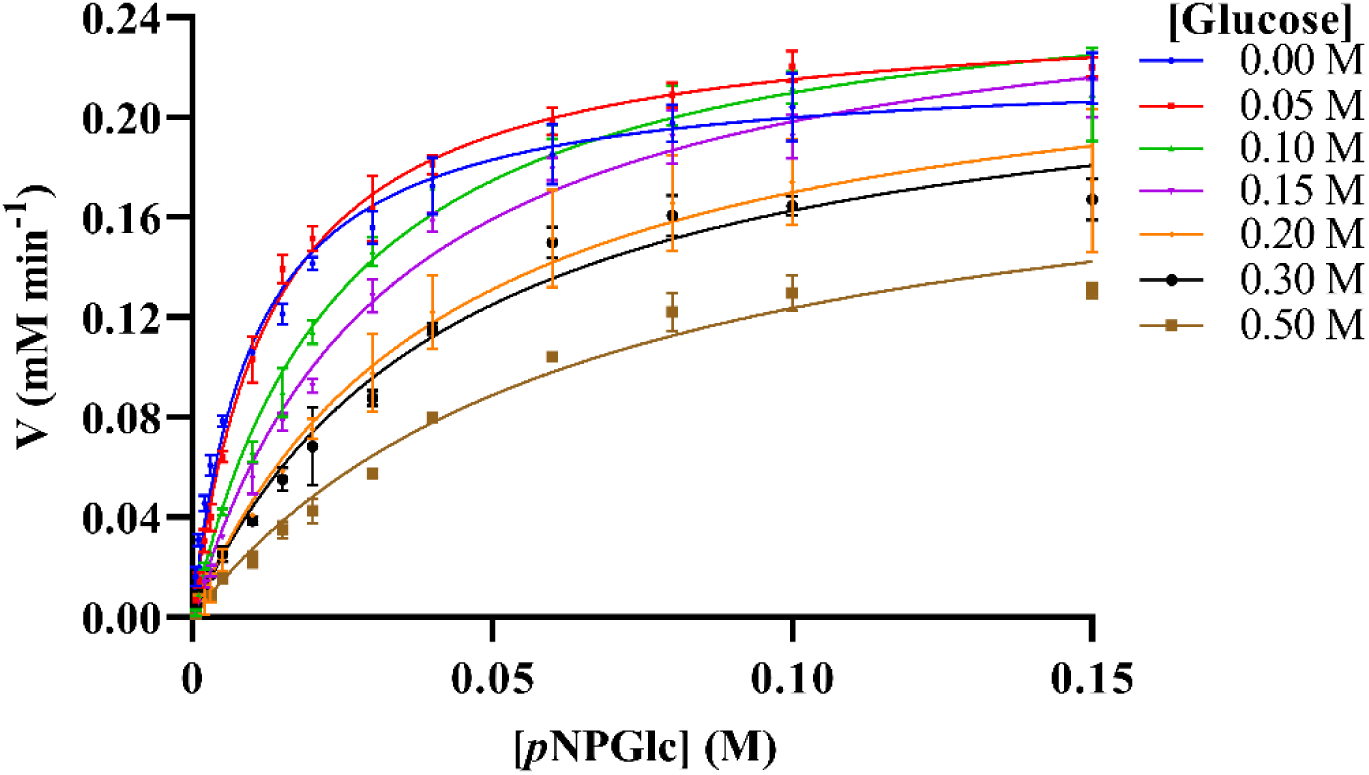
Non-linear regression (Michaelis-Menten Kinetics) with exogenous glucose (0-0.5 M) on substrate *p*NPGlc. The assay conditions are detailed in the methods section. All experiments were performed in triplicate and repeated at least twice. The standard deviations among the repeats were below 10 %. Complete kinetic data are presented in Table S1.

This biphasic kinetic behavior, characterized by initial stimulation at low glucose followed by progressive inhibition at higher concentrations, has been reported for other glycosidases and is often attributed to concentration-dependent modulation of enzyme conformational states^8, 18–21^. The observed increase in K_m_ suggests reduced apparent substrate affinity, while the decline in V_max_ at higher glucose concentrations suggests impaired catalytic turnover. Together, these features supported a mixed inhibition mechanism, as commonly observed in β-glucosidases and other glycosyl hydrolases^22^. Similar dual-phase responses, involving low-concentration activation followed by inhibition at elevated glucose levels, have been reported in both fungal and bacterial β-glucosidases^18, 21, 23, 24^. To further resolve the structural basis of this behavior, atomistic molecular dynamics simulations were performed to identify glucose interaction sites and their impact on BglB structural dynamics.

### Glucose-Induced Rigidification in BglB

To evaluate the structural impact of glucose on BglB, simulations were performed in the absence of glucose (C0) and in the presence of 0.09 M (C1), 0.3 M (C2), and 1.0 M (C3) glucose. These concentrations were selected based on kinetic behavior, corresponding to stimulation/retention (C1), partial inhibition (C2), and strong inhibition (C3) (Figure S1). In addition, a single glucose-bound state (C0.1) was analyzed to probe local interactions at the active site.

Across all conditions, BglB remained structurally stable over 1 µs, with global descriptors including RMSD, SASA, and secondary structure persistence (mean ± SEM across triplicates; Figure S2) showing no significant deviations from the energy-minimized structure, indicating preservation of the overall fold. In contrast, localized changes were observed in glucose-containing systems. A concentration-dependent rigidification trend was detected, particularly in residues lining the active-site entrance, most notably for residues in the gatekeeper region (Figure S2). These effects became more pronounced with increasing glucose concentration, consistent with enhanced glucose interactions in the gatekeeper region leading to localized stabilization of dynamics. Such glucose-induced constraints at the gatekeeper region may modulate BglB activity^7, 9, 25, 26^. At the same time, contributions from other regions cannot be excluded, as previous studies have implicated additional surface residues in glucose tolerance and inhibition^8, 20, 27–29^.

### BglB Inhibition is Mediated by Surface Secondary Binding Sites

In the C0.1 simulations, the single glucose molecule was found to exit the active site pocket within 100 ns, in all 3 replicates. Despite leaving the catalytic pocket, glucose formed multiple transient interactions with surface-exposed residues throughout the simulation (see Figure S3). This behavior is consistent with the experimentally observed non-competitive mode of inhibition and suggests the existence of additional sites that may facilitate glucose association. Motivated by these observations, we systematically investigated potential surface glucose-binding sites in the mixed-solvent simulations (C1-C3). Residue-wise glucose contact strength, 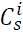, was quantified for each system (Figure 2a-c) and residues with 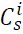 > 1.1, indicating contributions from at least two replicas, were identified as reproducible interaction sites (see Methods). Mapping these reproducible residues onto the protein surface revealed several discrete interaction patches. Spatially adjacent residues were subsequently grouped into eleven secondary binding sites (SBSs) (Figures 2g-h). Importantly, repeating the analysis using a hydrogen-bond distance criterion (3.8 Å) recovered highly similar sets of reproducible residues to those identified using the 7 Å distance cutoff, resulting in essentially the same SBS architecture with only minor differences in individual residue assignments (Figures S4 and S5). This demonstrates that SBS identification is robust to the choice of contact-distance criterion. Among the identified SBSs, two of the five highest-ranking clusters, SBS2 and SBS7, ranked according to the number of contributing residues and their cumulative contact strength (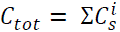) were located adjacent to the entrance of the active-site pocket (Figure 2d and Figure S6). These regions are physically connected to the aglycone site of the catalytic pocket through previously described gatekeeper residues and are therefore referred to here as gatekeeper-extended SBSs^30, 31^. The remaining SBSs were distributed across surface regions that are spatially separated from the active-site pocket (distal SBS), suggesting that glucose associates with BglB through multiple distinct surface interaction regions rather than a single dominant surface hotspot.

**Figure 2.**
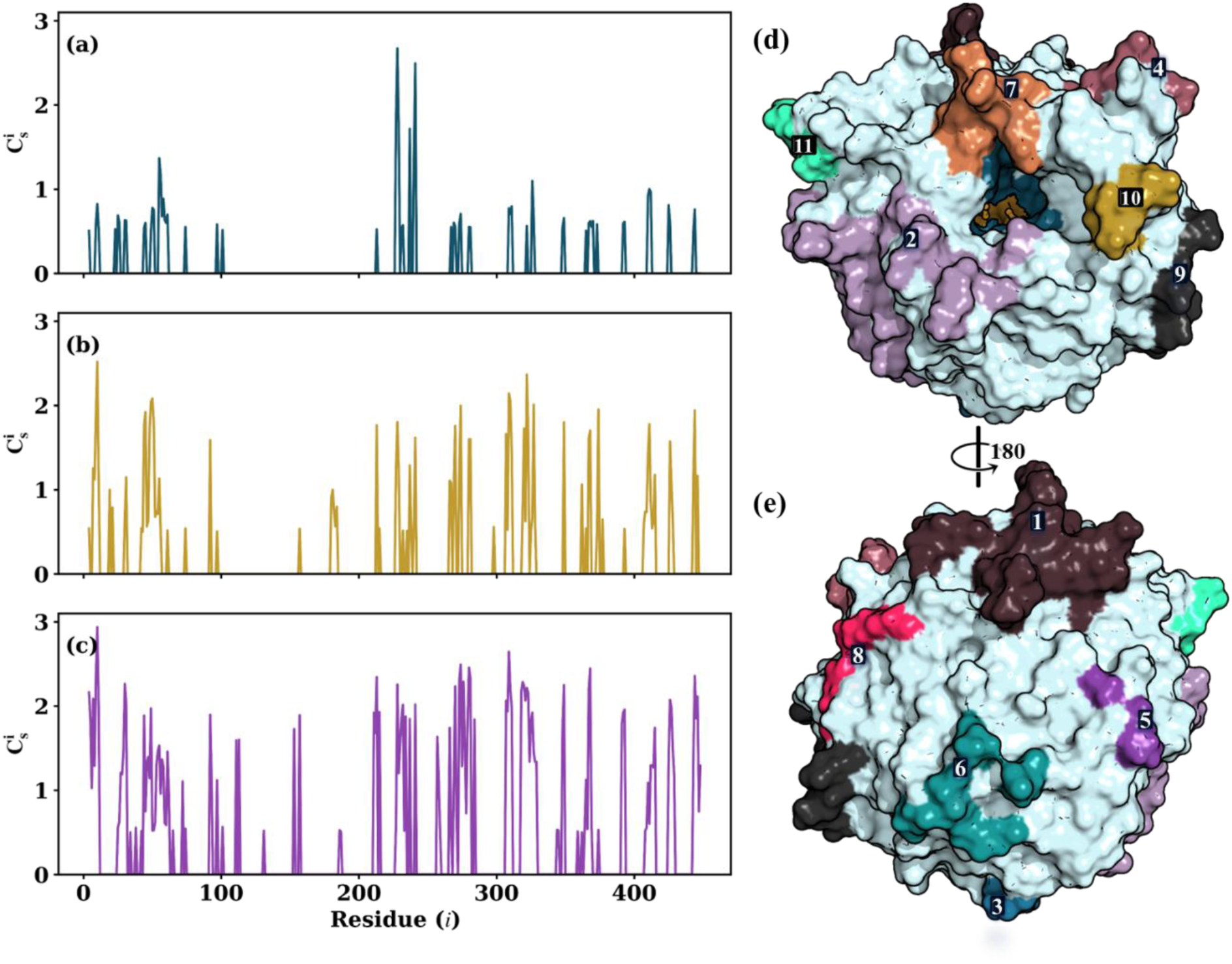
Residue-wise BglB-glucose contact strength at each concentration. The Heaviside-weighted consensus contact strength *C^i^* reflects the combined strength and reproducibility of residue (*i*)-glucose interactions from three independent replicas (see Methods). Panels (a-c) show the *C^i^* profiles for C1 (0.09 M), C2 (0.3 M), and C3 (1 M), respectively. Reproducible residues from *C^i^* were mapped onto the protein surface and grouped into spatial clusters, defined as secondary binding sites (SBSs), each shown in distinct colors (d-e). The protein surface is rendered in pale cyan, with the active-site pocket shown in teal, and catalytic residues Glu167 and Glu356 are highlighted as bright orange surface with sticks. Residue identities for each SBS are listed in Table S2, and the corresponding cumulative contact strength (*C_tot_*) of each SBS in Figure S6.

To determine whether these contact-defined SBSs also contributed to protein-glucose interaction energetics, cumulative protein-glucose interaction energies were estimated using the MM-GBSA approach (Δ*E*_int_, see Methods). Because multiple glucose molecules simultaneously interact with the protein surface in the cosolvent simulations, the calculated energies represent concentration-dependent cumulative interaction energies rather than the binding free energy of an individual glucose molecule. Accordingly, the MM-GBSA analysis can be viewed as a computational titration profile describing progressive glucose association with the BglB surface. The cumulative interaction energy became progressively more favorable with increasing glucose concentration, with Δ*E*_int_ changing from –44.6± 18.2 kJ mol^-1^ for the single-glucose system (C0.1) to –64.5 ± 11.2 kJ mol^-1^ (C1), –183.6 ± 17.7 kJ mol^-1^ (C2), and −660.4 ± 22.7 kJ mol^-1^ (C3). This trend is consistent with the concentration-dependent increases in contact probability (*p_ij_*) and residue contact strength (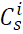), which resulted in a larger number of residues satisfying the reproducibility criterion, and consequently the identification of additional SBSs at higher glucose concentrations. Together, these results indicate that glucose association with BglB occurs through progressive recruitment of multiple spatially distinct surface interaction sites rather than through a single dominant interaction hotspot.

Further, to determine whether the contact-defined SBS residues also contribute directly to protein-glucose interaction energetics (Δ*E*_int_), residue-level interaction energies (IE) with glucose were evaluated. Residues identified from the contact probability analysis (*p_ij_* > 0.5) consistently exhibited progressively stronger energetic contributions with increasing glucose concentration (Figure 3a-c). Although a few residues contributed unfavorably, most SBS residues displayed increasingly favorable interaction energies (*IE*), demonstrating that residues with high glucose occupancy are also the principal energetic contributors to glucose interaction. Together, the consensus contact and energetic analyses suggest that the identified SBSs represent functionally relevant glucose-binding regions rather than merely recurrent interaction patches, contributing substantially to the cumulative protein–glucose interaction energy.

**Figure 3.**
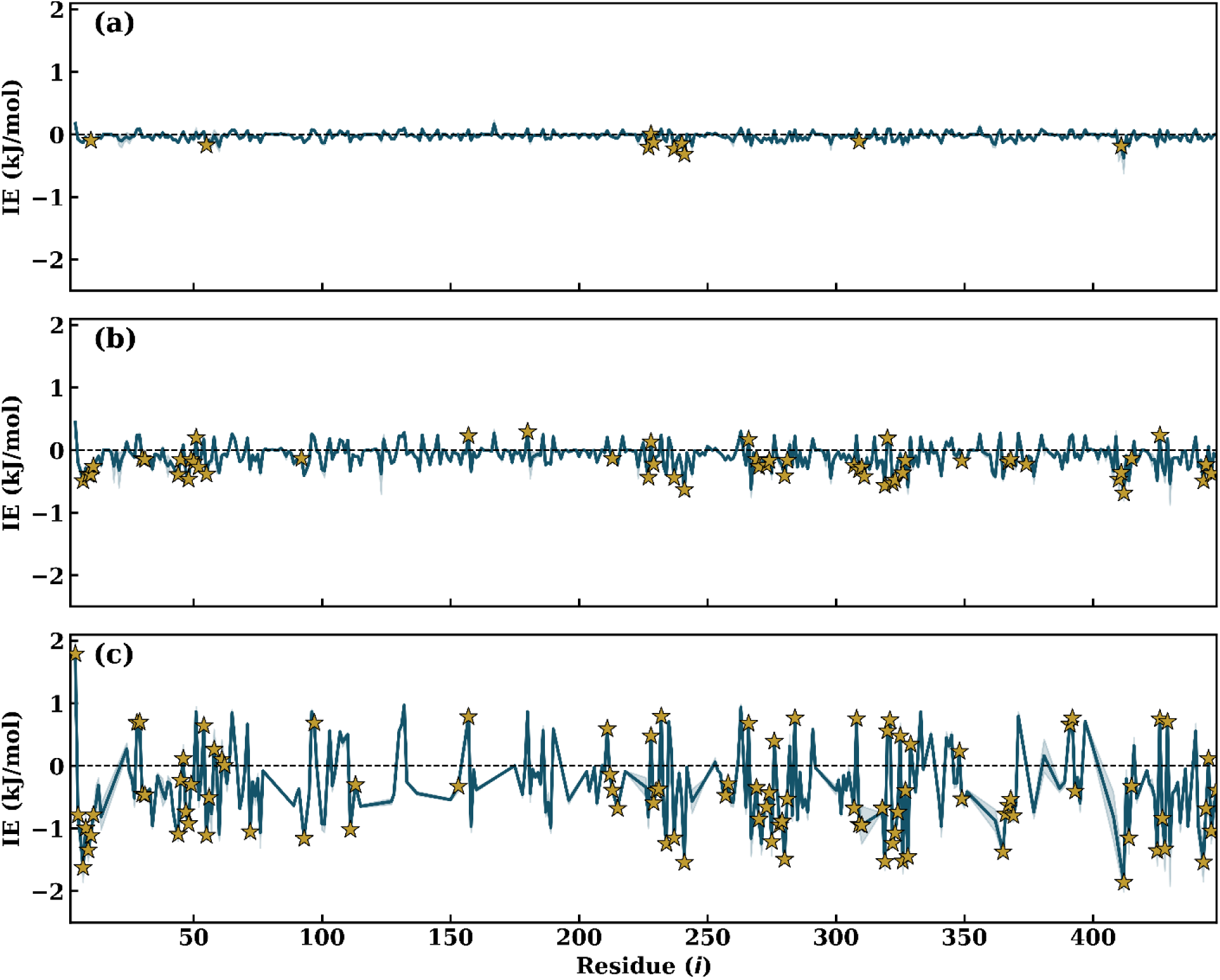
Per-residue energy decomposition (IE, mean ± SEM, across replicates) for WT at increasing glucose concentrations. (a) C1, 0.09 M (b) C2, 0.3 M and (c) C3, 1 M. Golden stars denote residues with a mean interaction probability (*p_ij_*) > 0.5, as determined from the contact probability analysis. The SEM is not readily discernible because of the minimal variation among the three independent replicas.

### SBS-Based Rational Mutant Design

To probe the structural and functional roles of the identified secondary binding sites (SBSs) and assess their influence on glucose binding, protein stability, and active-site regulation, a subset was selected for rational mutagenesis based on (i) proximity to the entrance of the active-site pocket (gatekeeper-associated), with potential roles in substrate access or product release; (ii) consistent engagement with glucose across concentrations C1-C3; and (iii) the size of the SBS, defined by the number of participating residues within each surface cluster. Accordingly, SBS2 and SBS7 were selected as gatekeeper-associated sites, while SBS4 was chosen based on its highest *C_tot_* at the lowest concentration and its ranking among the top sites across concentrations. In addition, SBS1, the largest distal cluster (non-gatekeeper associated), was included as it is spatially separated from the catalytic pocket and not directly involved in substrate binding, thereby enabling assessment of potential long-range effects on active-site regulation. At each selected SBS, non-conserved residues were substituted with the corresponding residues observed in glucose-tolerant GH1 homologs, with the goal of preserving overall structural integrity while modulating glucose interactions. For SBSs located near the active-site entrance (SBS2 and SBS7), only residues that were both non-conserved and functioned as gatekeepers were mutated to corresponding residues in the glucose tolerant homologue (Detailed in Table S3). In contrast, for distal SBSs not connected to the active-site pocket (SBS1 and SBS4), all non-conserved residues were mutated.

The resulting mutant sequences were modeled using AlphaFold3^32^, and the predicted structures were designated mSBS1, mSBS2, mSBS4, and mSBS7 according to their corresponding SBS index. Each mutant structure was aligned with the wild-type (WT) BglB to confirm preservation of the global fold and local conformational features (RMSD < 0.5 Å) and subsequently subjected to all-atom molecular dynamics simulations under the same conditions used for WT BglB. Simulations incorporating glucose were performed at 0.3 M to evaluate the impact of the mutations on glucose interactions.

### Dual Functional Role of SBS in BglB

*SBS Reshapes the Active-Site Pocket*: We first confirmed the structural integrity of all mSBS variants through molecular simulations. The global fold remained largely preserved, with RMSD, SASA, R_g_, and structural persistence (S^p^) profiles (mean ± SEM across triplicates, Figure S7) showing minimal deviation from the WT in both apo and glucose-bound states. RMSF profiles indicated that the mutated SBS regions were largely stable; however, residues exhibiting significant flexibility changes (ΔRMSF > 2 Å; Figure S7) were predominantly localized at gatekeeper regions, particularly in mSBS1, mSBS2, and mSBS4. Based on these observations, we next examined the impact of these mutations on active-site pocket dynamics. Interestingly, distal SBS mutations (mSBS1 and mSBS4) produced perturbations in pocket dynamics comparable to those observed for mutations proximal to the active-site entrance (mSBS2 and mSBS7) (Figure 4a-b), highlighting the presence of long-range coupling within the protein. As all mutants exhibited increased local SASA and decreased secondary structure persistence (S^p^) relative to the WT, indicating structural rearrangements at the active-site pocket (Figure 4a-b), we further analyzed the relative positioning of catalytic residues (Glu167 and Glu356 at subsite 0) with respect to each other and to the gatekeeper residues. This analysis revealed that the catalytic residues move farther apart from each other (Figure S8) while simultaneously approaching the gatekeeper residues, suggesting a broadening and concomitant shallowing of the active-site pocket.

**Figure 4.**
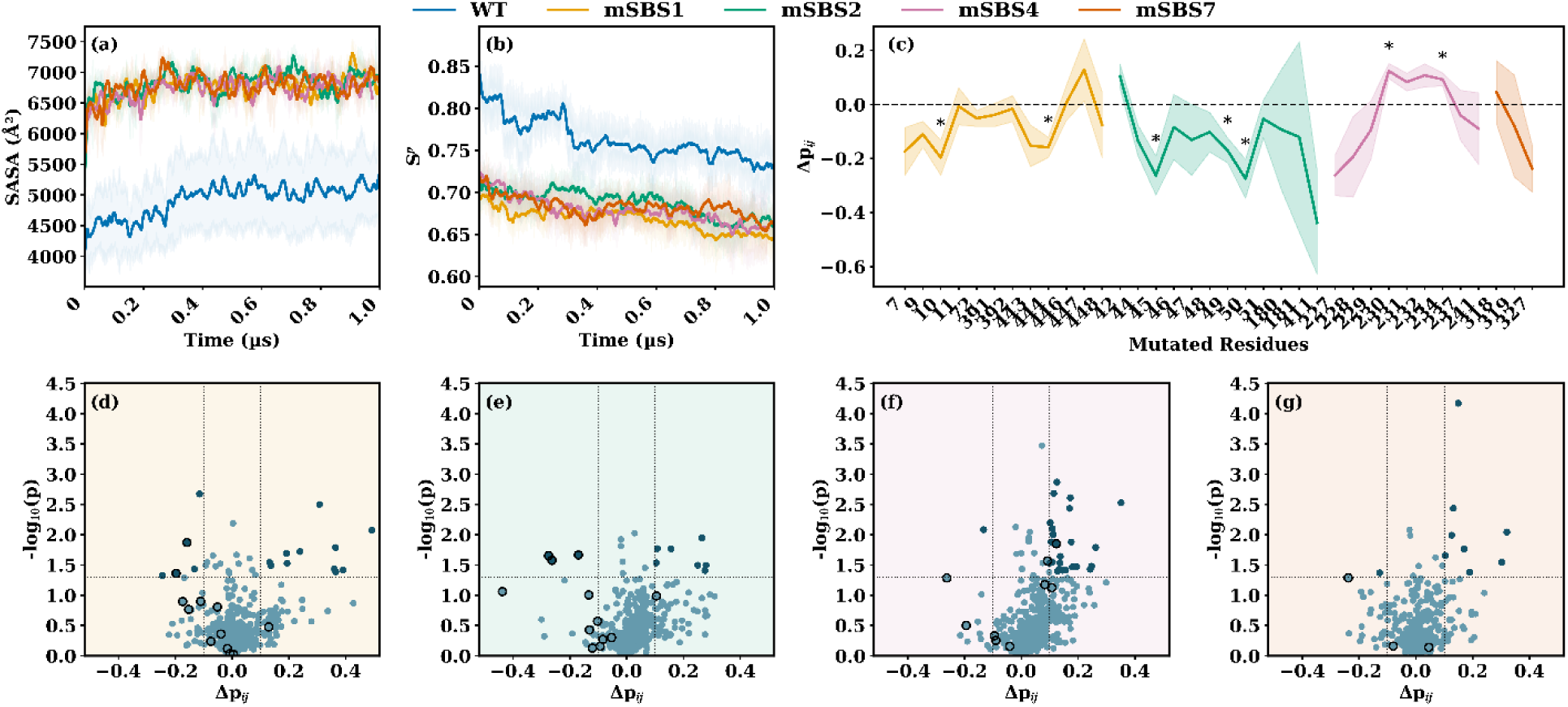
WT and mSBS structural and glucose interaction properties. (a-b) Time evolution of solvent-accessible surface area (SASA) and structural persistence (Sᵖ), respectively, for WT and mSBS systems in the presence of 0.3 M glucose. (c) Per-residue differences in mean contact strength (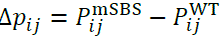) between mSBS and WT BglB for the respective mutated residues (X-axis) in mSBS1, mSBS2, mSBS4, and mSBS7. The dashed horizontal line at Δ*p_ij_*= 0 indicates no difference relative to WT, and residues marked with * denote statistically significant differences, based on Welch’s t-test. (d-g) Volcano plots showing residue-wise changes in contact strength (Δ*p_ij_*) for all protein residues in mSBS1, mSBS2, mSBS4, and mSBS7, respectively, relative to WT. Dark-colored points indicate residues satisfying the significance criteria (p < 0.05 and | Δ*p_ij_*| > 0.1). Across the first row, solid lines represent mean values over three independent replicas, and shaded regions denote the corresponding standard error of the mean (SEM). Colors consistently represent the mutant systems: mSBS1 (orange), mSBS2 (green), mSBS4 (purple), and mSBS7 (vermillion).

*SBS Contributes Directly to Glucose Binding*: SBS mutations were originally selected based on their expected role in glucose binding; to evaluate the impact of these mutations, we quantified glucose contact probability (*p_ij_*) at the mutated sites and across the protein. All SBS mutations generally reduced *p_ij_* at the targeted SBS (Figure 4c), supporting the assignment of these surface regions as glucose-binding sites in BglB. The magnitude of reduction varied among SBSs. In mSBS1, where 12 of the 16 SBS1 residues were mutated, 10 exhibited decreased *p_ij_*, with five showing reductions greater than 10%. In mSBS2, 11 of the 12 mutated residues showed reduced contacts, with decreases reaching ∼60% for some residues. Comparable, although less pronounced, reductions were observed for mSBS4 and mSBS7 (Figure 3c; Figure S9). Consistent with these observations, the mutated SBS regions displayed increased water residence times (Figure S10), indicating weaker glucose association and compensatory stabilization by water molecules. At the whole-protein level, however, only 9–25 of the 445 residues exhibited significant changes in glucose contact probability (Figure 4d–g), demonstrating that the effects of SBS mutations remained localized rather than globally disrupting glucose binding.

Despite the consistent reduction in glucose contact probability at the mutated SBSs, the overall Δ*E*_int_were only modestly affected. Comparison of cumulative protein-glucose interaction energies between WT and the four mSBS variants under identical glucose conditions (0.3 M; C2) showed only modest differences, with Δ*E*_int_ ranging from –156.33 ± 14.43 to –212.3 ± 20.46 kJ mol^-1^ compared with −183.6 ± 17.7 kJ mol⁻¹ for WT. Even the largest deviation (mSBS4; ΔΔ*E*_int_ = Δ*E*_mutant_ – Δ*E*_WT_; ≈ +27.1 kJ mol^-1^) remained within the same order of magnitude as the WT value, indicating that disruption of an individual SBS does not abolish overall glucose association with the BglB surface. However, per-residue energy decomposition revealed that several mutated residues underwent significant changes in interaction energy, with most exhibiting losses of favorable interactions and a smaller number displaying compensatory gains (Welch’s t-test, p < 0.05), whereas only a limited number of additional residues elsewhere on the protein were significantly affected (Figure 5). Across the mutated residues, the loss of residue-wise interaction energy, normalized relative to the WT, typically ranged from approximately 20–150%, with several residues showing losses exceeding 200% (Table S4). Thus, SBS mutations primarily redistribute local energetic contributions while preserving the overall cumulative protein-glucose interaction energy, indicating that glucose association is distributed across multiple surface binding sites rather than being dominated by any single SBS.

**Figure 5.**
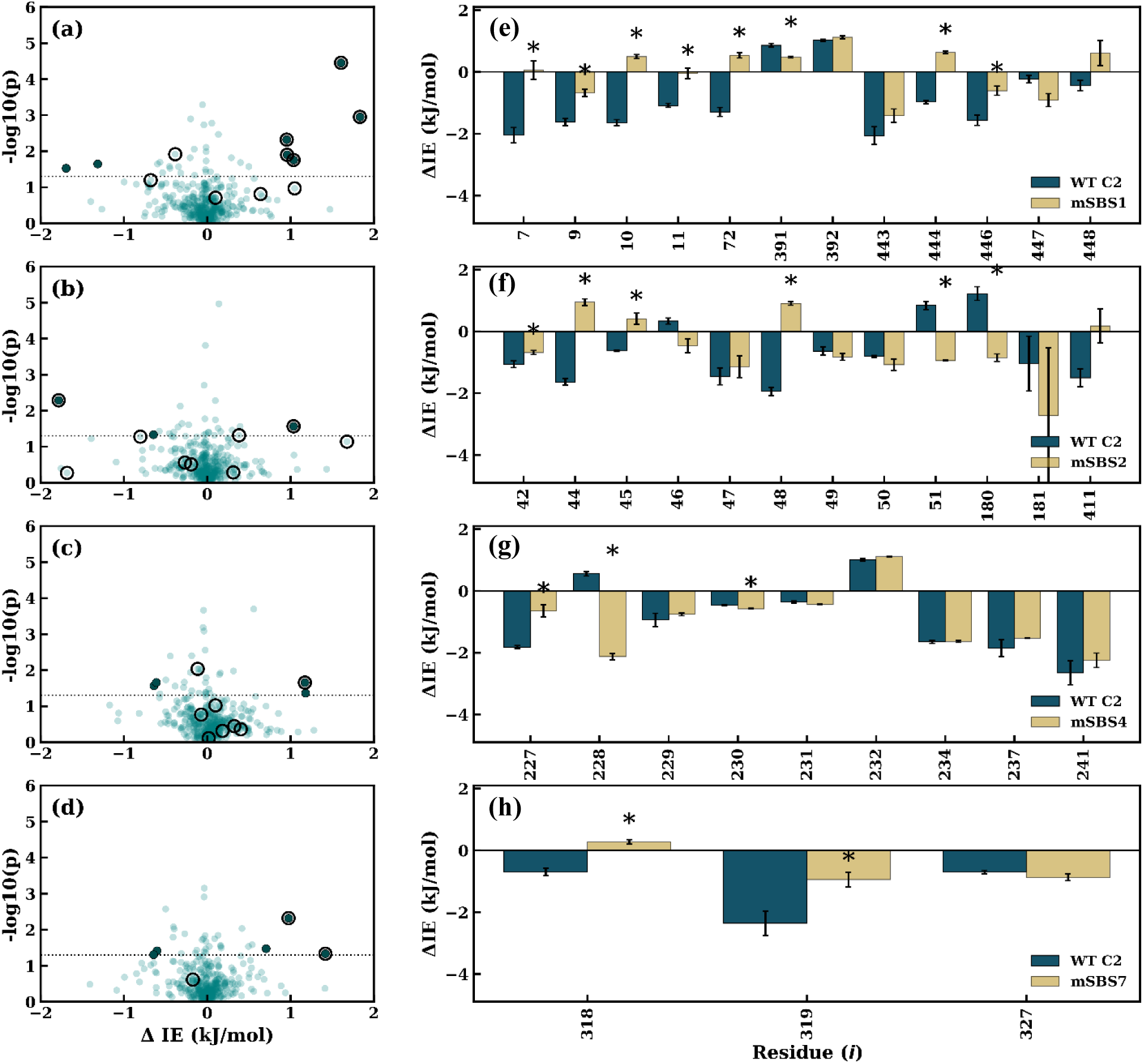
Comparison of per-residue energy contributions to protein-glucose interaction energy between WT C2 and the four mutant systems. Volcano plots showing the change in residue interaction energy, ΔIE (*IE*_mutant_ − *IE*_WT_; kJ mol⁻¹), plotted against statistical significance (−log₁₀ *p*). The horizontal dashed line denotes the significance threshold (*p* = 0.05); residues above this line exhibit statistically significant differences with the WT. Panels correspond to (a) mSBS1, (b) mSBS2, (c) mSBS4, and (d) mSBS7; in each, the respective mutated residues are circled. Bar plots compare the raw per-residue binding interaction energy contributions in WT C2 (teal) and the corresponding mutant (golden yellow) (mean ± SEM; *n* = 3 independent simulations) for only the residues mutated in each respective structure: (e) mSBS1, (f) mSBS2, (g) mSBS4, and (h) mSBS7, respectively. Asterisks (*) denote residues showing statistically significant differences (*p* < 0.05).

Despite modest effects on protein-glucose interaction energetics, the SBS mutants produced marked structural rearrangements at the active-site pocket. The unexpected observation prompted us to examine whether the mutations reorganized the intrinsic architecture of the protein itself. Receptor energy decomposition revealed substantially larger energetic perturbations than were apparent from cumulative protein-glucose interaction energies. Although the strongest changes occurred at the substituted SBS residues, significant energetic redistribution extended to neighboring and distant residues, particularly in mSBS1 and mSBS2 (Figure S11). These observations demonstrate that in the SBS mutations, internal energetic reorganization extends far beyond the mutational sites, providing evidence for long-range coupling between surface SBSs and the catalytic core. This observation prompted us to investigate whether SBS mutations alter communication pathways by inducing rewiring of the residue interaction network linking surface SBSs to the active-site pocket.

### SBS Mutations Redistribute Residue Communication Networks Coupling Surface Binding Sites to the Catalytic Core

Although the mSBS variants remained folded throughout the simulations and retained comparable overall glucose interaction energetics, residue-wise MM-GBSA decomposition revealed substantial redistribution of energetic contributions together with pronounced active-site rearrangements. These observations suggested that perturbations introduced at surface SBSs propagate beyond their immediate local environments and influence the catalytic core through long-range residue communication networks. To investigate the structural basis of this coupling, we employed protein structure network (PSN) analysis. Residue-level PSNs were constructed from inter-residue distance (*r*ᵢⱼ) and generalized correlation (*gcc*ᵢⱼ) matrices (see Methods for details).

The shortest communication paths between individual SBSs and the active-site pocket remained largely preserved across glucose concentrations and mutant systems (Figure S12), indicating that the overall communication routes are robust to mutation. However, inspection of the underlying residue-residue distance and generalized correlation matrices revealed substantial alterations in many of the interactions composing these pathways. Thus, although the topological routes remained largely unchanged, the residue interaction patterns supporting communication along these pathways were selectively reorganized.

To quantify this reorganization, Leiden-based community detection was applied to the residue interaction networks, and consensus communities obtained from triplicate simulations were aligned using the Hungarian algorithm (see Methods). Global similarity between WT and mutant networks was evaluated using complementary information-theoretic metrics: Normalized Mutual Information (NMI), Adjusted Rand Index (ARI), and Variation of Information (VI). NMI values (0.72–0.85) indicate that the overall modular organization of BglB remained largely preserved following mutation, with broadly similar groups of residues continuing to define corresponding communication communities. In contrast, lower ARI values (0.52–0.76), together with elevated VI values, demonstrate substantial redistribution of individual residue memberships within these communities. Because ARI evaluates pairwise residue co-assignment, even modest reassignment of residues between communities can substantially reduce ARI despite preservation of the overall modular framework. Collectively, these complementary metrics indicate that SBS mutations primarily induce selective local reorganization of residue interaction communities rather than wholesale disruption of the global communication architecture. Such behavior is consistent with emerging views that perturbations redistribute communication across pre-existing interaction networks rather than being transmitted through single dedicated pathways^33, 34^.

Community reorganization was most pronounced for mSBS1, which exhibited the lowest NMI and ARI values together with the highest VI, indicating the greatest deviation from WT communication organization, whereas mSBS4 and mSBS7 remained most similar to the WT network (Table 1). Notably, the extent of community reorganization closely paralleled the energetic redistribution observed in the MM-GBSA analyses. mSBS1 and mSBS2 displayed both pronounced energetic perturbation and extensive community reorganization, whereas mSBS4 and mSBS7 exhibited comparatively smaller energetic redistribution together with greater preservation of WT communication organization. These observations suggest that redistribution of residue interaction energetics is accompanied by corresponding reorganization of residue communication networks.

**Table 1.** Normalized Mutual Information (NMI) and Adjusted Rand Index (ARI) values for Hungarian-aligned consensus community partitions (derived from triplicate simulations) between wild-type (WT) and mutant (mSBS) systems under the 0.3 M glucose condition, quantifying global community similarity and pairwise residue assignment consistency. Additional comparisons across other glucose conditions are provided in Table S5.

| Pair | NMI | ARI | VI |
| --- | --- | --- | --- |
| WT, mSBS1 | 0.72 | 0.52 | 1.47 |
| WT, mSBS2 | 0.82 | 0.70 | 0.94 |
| WT, mSBS4 | 0.85 | 0.76 | 0.90 |
| WT, mSBS7 | 0.82 | 0.74 | 1.08 |

To determine how these network changes affected communication between the catalytic core and surface binding sites, we quantified community co-localization between the inner active-site pocket and each SBS (Figure S13). Community co-localization measures the probability that residues from two functional regions occupy the same communication module and therefore provides a localized measure of communication coupling complementary to the global partition similarity metrics. SBS mutations altered not only communication involving the mutated site itself but also coupling between distal SBS regions, indicating propagation of perturbations throughout the residue interaction network. For example, mutation of SBS1 reduced active-site co-localization with SBS1 itself (−20 residue pairs) while simultaneously increasing coupling involving SBS2 (+14 pairs) and SBS4 (+18 pairs). Similarly, mutation of SBS2 produced only modest local changes at SBS2 itself (+5 pairs) but induced the strongest increase in coupling between the catalytic pocket and the gatekeeper-associated SBS7 (+40 residue pairs). Likewise, mutation of SBS4 caused comparatively minor local effects (+9 pairs) while substantially increasing active-site coupling with SBS1 (+26 pairs) and SBS7 (+31 pairs). Thus, the largest communication changes frequently occurred in catalytic core–SBS communication pairs involving regions distal to the mutated SBS rather than at the mutated site itself. These observations demonstrate that perturbations/mutations introduced at one SBS redistribute communication coupling throughout the catalytic core–SBS interaction network, revealing that functional coupling between surface SBSs and the catalytic pocket arises from an interconnected communication architecture rather than from isolated site-specific pathways.

Collectively, these findings indicate that SBS mutations selectively reorganize residue interaction communities while preserving the overall communication architecture of BglB. Rather than remaining locally confined, mutation-induced perturbations are redistributed through interconnected residue communication networks that reshape how individual SBSs become coupled to the catalytic core.

Importantly, the present analyses should primarily be viewed as an attempt to understand how mutation-induced perturbations are redistributed through pre-existing residue interaction networks. Although the observed changes clearly demonstrate selective reorganization of communication pathways, they do not presently establish whether particular communication states are beneficial, compensatory, or detrimental for enzyme function^33–35^. Likewise, because the simulations were initiated from folded structures, the observed network alterations cannot be directly linked to folding efficiency or soluble expression. Instead, these analyses provide a mechanistic framework describing how distant surface perturbations influence active-site organization through redistribution of residue communication patterns^34, 36, 37^. Determining which communication architectures remain compatible with productive folding and function therefore requires experimental interrogation of selected mutants.

### Glucose Tolerance in mSBS7

To assess how mutation-induced redistribution of residue communication networks produces experimentally observable phenotypic consequences, representative SBS mutants were expressed and biochemically characterized. His-tagged WT and mutant constructs were expressed in *E. coli*. Among the four designed variants, only mSBS7 was obtained as a soluble, purified protein. In contrast, mSBS1 and mSBS4 were recovered predominantly in the insoluble fraction, whereas mSBS2 co-purified extensively with host proteins (Figure S14), preventing further biochemical characterization. These observations indicate that substitutions at SBS1, SBS2, and SBS4 adversely affected recombinant protein production, most likely through impaired folding, reduced solubility, or decreased structural stability during biosynthesis. Circular dichroism spectroscopy further showed that mSBS1 exhibited a marked reduction in secondary structural content, whereas mSBS7 closely resembled the WT spectrum with only minor differences (Figure S15), indicating preservation of the native structural framework in mSBS7 but compromised structural integrity in mSBS1.

Because mSBS7 was the only mutant that could be purified as a soluble protein while preserving a WT-like secondary structure profile, it was selected for detailed biochemical characterization. Temperature and pH profiling showed that the mutant retained optima comparable to those of the WT, indicating preservation of the catalytic framework. Under identical assay conditions, mSBS7 retained approximately 97% of the WT catalytic efficiency (Figure 6). In the absence of exogenous glucose, substrate affinity improved substantially, with *K*_m_ decreasing from 10.16 mM to 6.3 mM (≈38% reduction). Furthermore, glucose-mediated inhibition was alleviated, as evidenced by an increase in the inhibition constant (*K*_i_) from 95 mM to 124 mM (≈31% increase). Collectively, these results demonstrate that perturbation of the gatekeeper-associated SBS7 enhances glucose tolerance while preserving catalytic efficiency.

**Figure 6.**
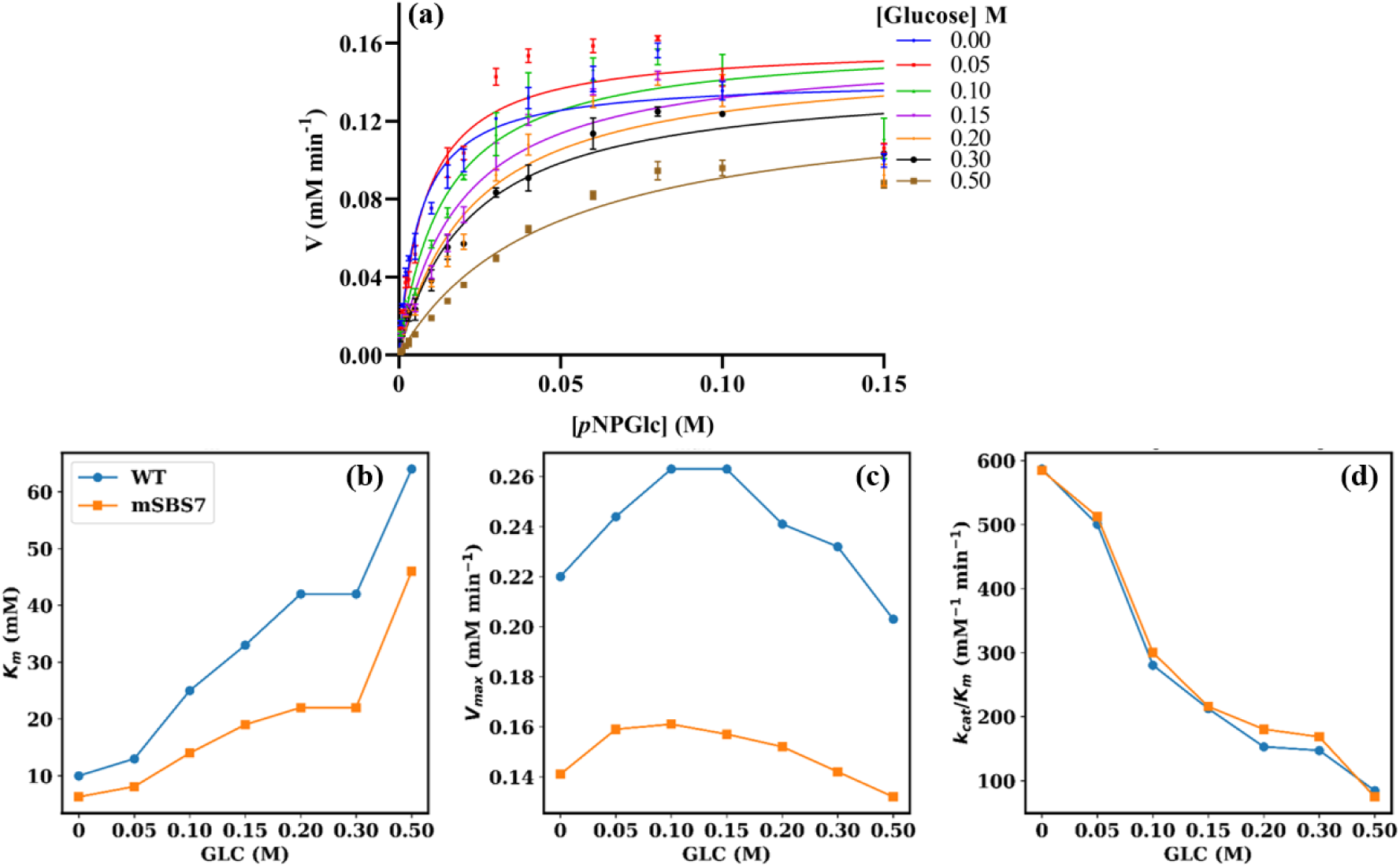
Michaelis-Menten kinetics of the mSBS7 mutant in the presence of exogenous glucose (0-0.5 M) using *p*NPGlc as the substrate. (a). Relative changes in K_m_, V_max,_ and catalytic efficiency (k_cat_/K_m_) for WT BglB and mSBS7 are shown in panels (b-d), normalized to the WT in the absence of glucose (0 M), which was used as the reference condition. To facilitate comparison of the combined effects of glucose concentration and mSBS7 mutation on enzyme performance.

At first glance, the insolubility of mSBS1 and mSBS4 together with the unsuccessful purification of mSBS2 appears inconsistent with the simulation results which indicate that the folded mutant structures remained stable over the simulated timescale. This apparent discrepancy reflects the different biological processes being examined. In the absence of experimentally determined mutant structures, the simulations were initiated from pre-folded models that exploit available databases to predict plausible end-state stability while ignoring the folding pathways^38^. In contrast, recombinant expression requires successful co-translational folding, maturation, and maintenance of solubility before a folded protein can accumulate. Consequently, mutations may perturb folding or promote aggregation during biosynthesis without destabilizing the folded state sampled during simulation. Thus, the computational analyses indicate that the folded conformations are intrinsically stable, whereas the experimental observations demonstrate that several SBS mutations impair the ability of the protein to attain those conformations *in cellulo*.

Taken together, these findings indicate that SBS7, located adjacent to the active-site entrance, contributes to glucose-mediated product inhibition in WT BglB. Glucose interaction at this gatekeeper-associated SBS is proposed to promote local crowding at the substrate-entry pathway, thereby hindering substrate access and perturbing substrate positioning at the gatekeeper region, which is known to regulate substrate entry^31,39^. In contrast, perturbation of the distal SBS1 resulted in loss of soluble protein production despite the computationally predicted stability of its folded state, indicating that this SBS contributes not only to glucose binding but also to maintaining the structural framework required for productive folding and enzyme function.

Integrating computational and experimental observations, we propose that SBSs collectively serve dual roles in BglB. First, they directly contribute to glucose association, with gatekeeper-associated SBSs modulating glucose-mediated inhibition by influencing substrate access near the active-site entrance.

Second, SBSs participate in long-range residue communication networks that couple surface perturbations to active-site organization. Mutations at individual SBSs redistribute energetic interactions and alter communication coupling throughout the protein, thereby modifying active-site organization despite preservation of the global fold. Such behavior is consistent with emerging views that functional regulation can arise through redistribution of communication networks across pre-existing interaction architectures rather than through transmission through single dedicated pathways^33, 34^. In this context, distal SBSs such as SBS1 may contribute not only to glucose binding but also to maintaining communication architectures compatible with productive folding and enzyme function. Together, these findings extend previous studies that associated surface or gatekeeper residues with glucose tolerance primarily through mutational or docking analyses^8, 20, 21, 27–29^, by providing a mechanistic framework linking surface glucose binding to active-site remodeling, energetic redistribution, and residue interaction network rewiring.

## Conclusions

This study provides integrated computational and experimental evidence supporting the existence and functional significance of secondary glucose-binding sites (SBSs) in the GH1 β-glucosidase BglB. Rather than functioning as passive surface interaction patches, these regions emerge as regulatory elements that couple glucose association to enzyme structure and function. Our results indicate that glucose interacts with BglB through multiple surface-exposed SBSs in a concentration-dependent manner, and that these interactions are accompanied by localized and long-range structural, energetic, and residue interaction network rearrangements that remodel the active-site environment.

Targeted mutations of representative SBSs revealed that their functional contributions depend strongly on their structural context. Mutation of the gatekeeper-associated SBS7 reduced glucose-mediated product inhibition while preserving catalytic activity, identifying this region as a determinant of glucose tolerance and substrate access. In contrast, mutation of distal SBS1 impaired soluble protein production despite preservation of stable folded states during simulation, suggesting that distal SBSs may also contribute to maintaining communication architectures compatible with productive folding and enzyme function. Together, these findings provide a mechanistic framework linking surface glucose association to active-site regulation through distributed residue communication networks within the protein.

Although experimental validation was limited to the soluble mSBS7 variant, the close agreement between computational predictions and biochemical measurements supports the proposed mechanistic model and motivates further investigation of additional SBSs. More broadly, our findings suggest that the functional importance of surface residues is shaped not only by their proximity to the catalytic center or sequence conservation but also by their participation in long-range communication networks. This perspective expands our understanding of product inhibition in β-glucosidases and identifies secondary glucose-binding sites as promising targets for engineering glucose-tolerant glycoside hydrolases with improved catalytic performance.

## Methods

### Cloning, Expression, and Purification of BglB

The gene encoding β-glucosidase from *Paenibacillus polymyxa* was constructed as previously described^30^. The construct was then transformed into *Escherichia coli* BL21(DE3) (Life Technologies, La Jolla, CA) and expressed. Following the expression, the cells were centrifuged at 4000 ×g for 10 minutes at 4 °C, and the resulting pellet was stored at −20 °C until purification. The protein was purified according to the earlier established protocol, and its purity was confirmed using 10 % SDS-PAGE^30^.

### Kinetics and Inhibition Assays

The effect of glucose on BglB activity was measured by varying the substrate *p*-Nitrophenyl-β-D-glucopyranoside (*p*NPGlc) concentration between 0.005 and 0.15 M, while maintaining constant glucose concentrations. The glucose inhibition constant, K_i_, of BglB was determined under optimal conditions of 42°C and pH 6.5, as reported earlier^30^. All assays were conducted in triplicate, using *p*NPGlc kinetic constants at varying substrate concentrations (0.005 M to 0.150 M). Glucose concentrations ranged from 0 M, 0.10 M, 0.15 M, 0.20 M, 0.30 M, 0.40 M, and 0.50 M. The experimental data were analyzed by fitting the results to the Michaelis-Menten equation using GraphPad PRISM version 9.0 (GraphPad Software, La Jolla, CA).

### Circular Dichroism (CD)

Circular dichroism (CD) spectra of all enzymes were recorded using a JASCO J-815 spectropolarimeter (Easton, MD) at room temperature in a 1 cm path-length quartz cuvette (Starna, Atascadero, CA). Data were collected with a bandwidth of 1 nm, a step resolution of 0.1 nm, a scan speed of 50 nm min⁻¹, and a response time of 0.25 s. The far-UV spectra were recorded over the wavelength range of 190–250 nm, while the near-UV spectra were acquired from 245 to 360 nm. Spectra were background-corrected by subtracting the appropriate buffer blank and subsequently smoothed. Enzyme samples (2.5 μM in 50 mM sodium phosphate buffer, pH 7.0) were used for both far-UV and near-UV measurements. High-tension (HT) values were confirmed to be ≤600 V for all spectra. Raw ellipticity values recorded in millidegrees (mdeg) were converted directly to molar ellipticity, [θ] (deg·cm²·dmol⁻¹), according to:

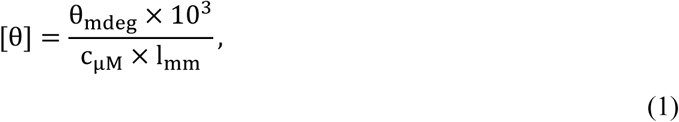

where *θ*_mdeg_ is the observed ellipticity in millidegrees, *c_μ_*_M_ is the enzyme concentration in μM, and *l*_mm_ is the cuvette path length in mm (1 mm in this study).

### Molecular Dynamics (MD) Simulation

Simulations of the X-ray crystallographic structure of BglB with a glucose-bound structure (PDB ID: 2O9T) were conducted to investigate interactions between BglB and its product (C0.1) at the active site pocket, similar to those observed with substrates^30^. Various systems were prepared with the protein in different glucose concentrations (C1, C2, and C3), while a previously reported control system in pure water was designated as C0. The number of glucose molecules required for cosolvent simulations was determined using the Held and Nagan method^40^. Additionally, the minimum distance of any atom from the box edge was maintained at 1.5 nm within a volume of 680 nm^3^. All simulations utilized GROMACS^41^, employing the CHARMM36m force field for protein, the TIP3P water model modified for CHARMM, and the CGenFF online server for glucose topology preparation^42–44^. All simulations were performed at the experimentally determined optimal temperature of 315 K, using a V-rescale thermostat and Parrinello-Rahman barostat. A detailed simulation protocol was reported previously^30^.

Following the identification of surface glucose binding sites (SBS), residues within these sites were mutated to corresponding residues found in glucose-tolerant GH1 β-glucosidase, based on structure-guided sequence alignment. The mutant BglB sequences were modelled using AlphaFold^45^. Each SBS mutant-SBS1, SBS2, SBS4, and SBS7-was subjected to MD simulations under conditions analogous to those of wild-type BglB to evaluate the effect of mutations on protein dynamics. A summary of the system details can be found in Table S6. All simulations were performed in triplicate, and further analyses were done and reported based on these independent replicas.

### Secondary Structure Persistence

The structural persistence (S^P^) of the backbone atoms in proteins was assessed for each trajectory across various conditions. S^P^ offers valuable insights into conformational changes among different protein states^46, 47^. The S^P^ value ranges from 0 to 1, with 1 indicating no deviation from the reference structure and lower values indicating greater deviations. The S^P^ was calculated using the following formula^47, 48^:

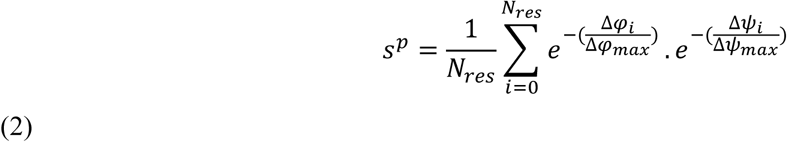

Here, *N*_res_ is the total number of residues in the reference protein structure, Δφ_i_ and Δψ_i_ represent the changes in the torsional angles (φ and ψ) for the i^th^ residue, and Δφ_max_ and Δψ_max_ denote the maximum possible torsional angle deviations on the Ramachandran plot.

### Surface Binding Sites (SBS)

Glucose was treated as the cosolvent. Contact strength for each residue *i* was evaluated across three independent trajectories. For each residue *i* and trajectory *j*, the fractional occupancy *p_ij_* is estimated as the fraction of simulation frames in which the residue remained within a 7 Å cutoff distance of any glucose molecule. Thus, *p_ij_* ∈ [0,1] represents the empirical probability of interaction in trajectory *j*.

To remove transient interactions, a Heaviside step function was applied to impose a persistence threshold:

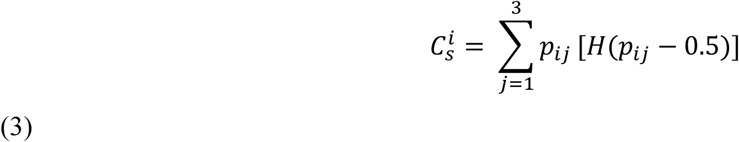

where *H*(*x*) is the Heaviside step function, defined as *H*(*x*) = 1 for *x* ≥ 0 and *H*(*x*) = 0 otherwise. A threshold of 0.5 was chosen to retain interactions present in at least 50% of the simulation frames, thereby filtering out transient contacts. This formulation ensures that only interactions with substantial occupancy (≥ 0.5) contribute to the final score, while weak or transient contacts are excluded. The resulting 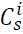 quantifies the reproducibility of residue-glucose interactions across independent simulations, ranging from 0 (no significant interaction in any trajectory) to 3 (consistent interaction across all the 3 simulation trajectories). Residues with 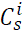 > 1.1 were selected for downstream identification of stable surface binding sites. The primary analysis employed a 7 Å residue–glucose distance cutoff, thereby capturing both direct hydrogen-bonding and proximal non-bonded interactions associated with surface binding. As a robustness analysis, the complete SBS identification workflow was independently repeated using a hydrogen-bond distance cutoff (3.8 Å), and the resulting SBS assignments were subsequently compared with those obtained using the 7 Å criteria.

### Protein-Glucose Interaction Energy and Receptor Energy Decomposition

Protein-glucose interaction energies and residue-level energy decompositions were calculated using the Molecular Mechanics Generalized Born Surface Area (MM-GBSA) approach implemented in gmx-MMPBSA^49^. MM-GBSA provides an efficient framework for estimating relative interaction energetics between receptor and putative ligands (here cosolvent glucose) by combining molecular mechanics energies with an implicit solvent model. During the calculations, explicit solvent molecules were removed from each trajectory frame and replaced by a Generalized Born continuum solvent representation.

Cumulative protein-glucose interaction energies Δ*E*_int_ were calculated using the single-trajectory protocol, in which the receptor, ligand and complex energy terms were extracted from the same simulation trajectories. For the C0.1 system, the ligand corresponded to the single glucose molecule initially placed in the catalytic pocket. For the glucose cosolvent simulations (C1–C3), all glucose molecules present in the simulation box were collectively treated as the ligand ensemble. Accordingly, the resulting Δ*E*_int_ values represent cumulative protein-glucose interaction energies rather than the binding affinity of an individual glucose molecule. The cumulative protein-glucose interaction energy was computed as

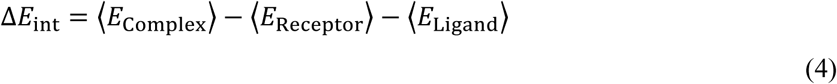

and was estimated within the MM-GBSA framework as

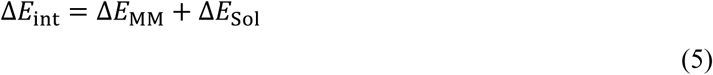

where *E*_Complex_, *E*_Receptor_, and *E*_Ligand_ represent the effective MM-GBSA energies of the protein-glucose complex, the isolated protein, and glucose ensemble, respectively. The molecular mechanics energy (*ΔE*_MM_) comprises bonded, electrostatic, and van der Waals interactions, and the solvation energy term (*ΔE*_Sol_) consists of polar (Generalized Born) and nonpolar components. Entropic contributions were not included; therefore, the reported values should be interpreted as relative interaction energies dominated by enthalpic contributions rather than rigorous thermodynamic free energies.

To identify energetic contributions at residue resolution, residue-wise MM-GBSA decomposition was performed. Two complementary decompositions were analyzed. Protein-glucose interaction energy decomposition quantified the contribution of individual residues to cumulative protein-glucose interactions. whereas receptor interaction energy decomposition evaluated residue interaction energies (IE) within the protein using the receptor energy term extracted from the same single-trajectory calculations. The latter analysis was used to determine how SBS mutations redistribute the intrinsic energetic organization and residue-residue interaction landscape of BglB independently of direct protein-glucose interactions.

### Dynamic Cross-Correlation Analysis

After aligning each trajectory on the C_α_ atoms to remove overall translational and rotational motions, residue-level descriptors were extracted for correlation analysis. Three independent replicas were analyzed for each system, and trajectory parsing, frame selection, and replica handling were performed using the MDigest^50^. Dynamic cross-correlations (*dcc_ij_*) between residues *i* and *j* were derived from cartesian atomic displacements, and torsional angle displacement matrices were calculated using MDigest^50^. For each replica, correlation matrices derived from the atomic displacement matrix and torsional angle matrix were averaged to obtain a residue-wise *N* × *N* cross-correlation matrix. Replica-specific matrices were subsequently averaged to generate the final consensus correlation matrix for each system.

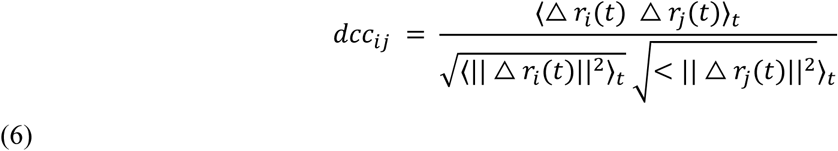

where Δ*r*_i_ (t) denotes the atomic or torsional fluctuation of residue *i* at time *t*. To extend beyond linear couplings, generalized correlation coefficients (*gcc_ij_*) were evaluated from mutual information (MI) estimated from a previously described Gaussian estimator^50, 51^.

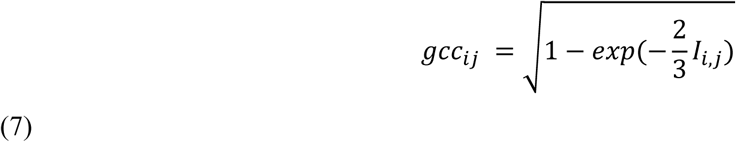

### Protein Structure Network

Using the computed distance and correlation matrices, residue-level protein structure networks were constructed using NetworkX^52^. In graph theory, a network is formally represented as a graph G = (V, E), where V denotes the set of vertices (nodes) and E the set of edges (connections between nodes). In the present study, each protein was represented as a residue-level PSN, with each vertex corresponding to the C_α_ atom of an amino acid residue. Edges were defined purely by geometric criteria: two residues *i* and *j* were connected by an edge if the Euclidean distance, *E*, between their C_α_ atoms was within 7 Å in the simulation ensemble^53, 54^.

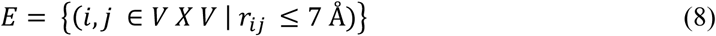

where *r_ij_* is the average distance between residues *i* and *j*.

Although the existence of an edge was determined only by geometry, generalized correlation coefficients (*gcc_ij_*) were computed to quantify the strength of dynamical coupling between residues. In the PSN representation, *gcc_ij_* values were not used to define connectivity but were instead mapped to the visual encoding of the network, with *gcc_ij_* encoded as edge color intensity. This allows both the structural connectivity (geometry) and the strength of dynamical correlation (*gcc_ij_*) to be captured within the same representation. The resulting PSN was thus an undirected graph encoding the spatial proximity of residues through edges, while simultaneously illustrating their dynamical correlations through edge coloring.

### Community Detection and Alignment

In a protein structure network (PSN), a community represents a group of residues that are more densely connected (*r_ij_* ≤ 7 Å) and dynamically coupled (*gcc_ij_* ≥ 0.2) with each other than with the rest of the protein, often corresponding to functional or structural subdomains^53–55^. The *gcc_ij_* ≥ 0.2 threshold was used to exclude weak dynamic correlations and retain only sufficiently strong residue couplings. To identify these communities *C* = (*C*_1_, *C*_2_, …, *C_k_*), we applied the Leiden modularity maximization algorithm, which partitions the PSN by optimizing intra-community connectivity while maintaining stability and high modularity^56^. Edges were weighted using generalized correlation coefficients (*gcc_ij_*), capturing dynamic coupling between residues. To ensure robustness, community detection was performed across multiple resolution parameters and random initializations, and the most stable partition (resolution = 1.0) was selected based on convergence across independent runs. Community assignments obtained from the three replica-specific networks were subsequently compared, and consensus communities were constructed from reproducible residue-grouping patterns shared across triplicates, thereby minimizing trajectory-specific fluctuations and capturing stable dynamical organization.

To compare community organization across wild-type (A) and mutant (B) systems, a one-to-one correspondence between communities was established using the Hungarian alignment algorithm, which determines the optimal mapping between partitions by maximizing overall similarity^57,58^. Global similarity between partitions was evaluated using a contingency-based partition comparison framework. A contingency matrix N = [*n_ij_*] was constructed, where rows correspond to communities in partition A (WT) and columns correspond to communities in partition B (mutant). Each entry *n_ij_* denotes the number of residues simultaneously assigned to community *i* in partition A and community *j* in partition B. From this matrix, a joint probability distribution was defined as *P*(*i*, *j*) = *n_ij_*/*N*, where *N* is the total number of residues. The corresponding marginal probabilities *P*(*i*) and *P*(*j*) were obtained from the row and column sums of the contingency matrix. This formulation captures residue-level reassignment of community membership across conditions and follows standard definitions of joint and marginal probability distributions in information theory^59^.

Normalized Mutual Information (NMI) was computed to quantify the shared information between partitions:

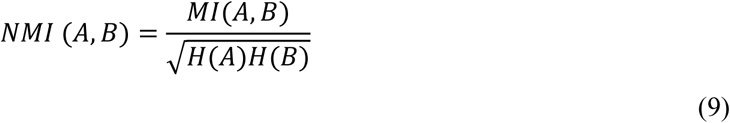

where,

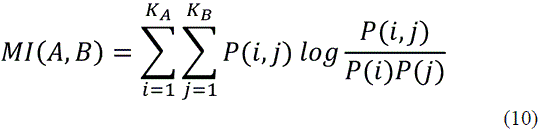

Here, *K_A_* and *K_B_* denote the number of communities in partitions A and B, respectively, and *H*(*A*) and *H*(*B*) represent the corresponding Shannon entropies^60, 61^. NMI measures the deviation of observed community overlap from statistical independence and thus reflects the extent to which community identities are preserved across systems.

In parallel, the Adjusted Rand Index (ARI) was used to assess pairwise consistency of residue assignments^62, 63^. Based on the contingency matrix *N* = [*n_ij_*], ARI^64, 65^ was defined as:

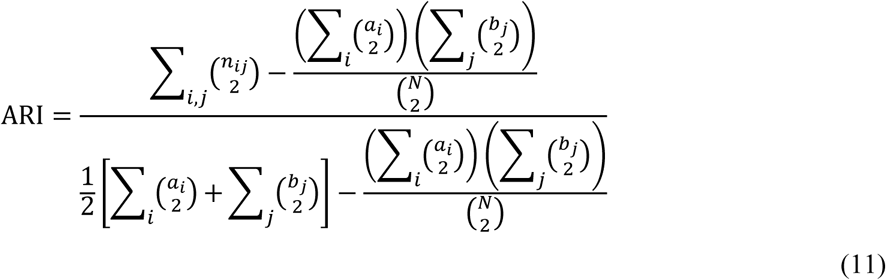

where 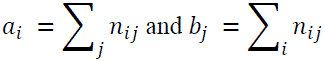 represents the sizes of community *i* in partition A and community *j* in partition B, respectively. ARI evaluates the agreement between all residue pairs that are either grouped together or assigned to different communities in two partitions, corrected for random expectation. Variation of Information (VI) was additionally calculated using the Shannon entropies of partitions as^59, 61^:

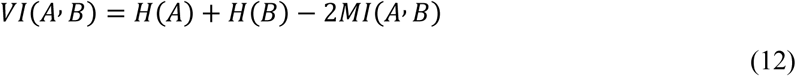

Unlike NMI, which measures shared information, VI quantifies the amount of information lost and gained when transitioning between partitions and therefore provides a direct measure of community reorganization^66^. Lower VI values indicate more similar community organization, whereas larger values reflect increased residue reassignment and network restructuring. In addition to these global measures, we defined functional coupling between predefined residue sets within each system as a pairwise co-localization metric:

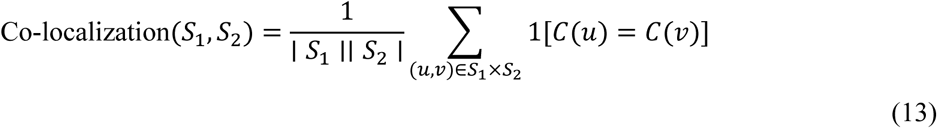

where the indicator function equals 1 if residues *u* and *v* belong to the same community and 0 otherwise. This represents the probability that a randomly selected residue pair from the two sets is assigned to the same community and provides a local measure of functional organization, complementary to global partition similarity metrics. High agreement across these complementary metrics indicates preserved community organization, whereas weak overlaps highlight mutation– or concentration-sensitive regions.

## Author Contributions

Conceptualization and study design: S.S., S.D., and N.S. Computational investigation and data analysis: S.S. Mutant design: S.S., S.D., and N.S. Biochemical assays and experimental data analysis: Sa.S. and S.S. Data interpretation: S.S., S.D., and N.S. Writing original draft: S.S. review and editing: S.S., S.D., and N.S. Supervision: S.D. and N.S.

## Acknowledgements

S.S. would like to thank Mr. Abel Francis Xavier for his assistance with the code for water residence calculations. N.S. acknowledges computational resources obtained through the “PARAM Rudra” (SN Bose National Centre for Basic Sciences), implemented by C-DAC and supported by the Ministry of Electronics and Information Technology (MeitY), support from ANRF via the ARG grant (ANRF/ARF/2025/000861/CS). S.D. acknowledges the Anusandhan National Research Foundation (ANRF), Government of India, CRG/2023/002111, and Department of Biotechnology (DBT), Government of India, BT/PR47801/BCE/8/1812/2023, IISER Kolkata Academic Research Fund, and the Center for Climate and Environmental Studies (CCES) for computational resources.

## Data and Code Availability Statement

The data that support the findings of this study are available in the Supporting Information of this article. The analysis scripts implementing the computational workflow developed in this study will be made publicly available upon publication through GitHub.

## Conflict of Interest

The authors declare that they have no conflicts of interest with the contents of this article

## Notes

### Competing Interest Statement

The authors have declared no competing interest.

