## Supplementary Information for "Surface Binding Sites Orchestrate Long-Range Control of Active-Site Dynamics and Glucose Tolerance in β-glucosidase, BglB"

**Contents**

**Figure S1.** Changes in relative activity of BglB with increasing concentrations of exogenous glucose (0-1M)....................................................................................................................................6

**Figure S2.** "Global" Structural characteristics of BglB in different glucose concentrations. Backbone RMSD timeline, C-alpha RMSF (highlighted gatekeeper residues), secondary structure persistence and SASA timeline for BglB. In each plot, lines represent mean values across three independent replica and shaded lines represent standard error of mean (SEM). ........................................................................7

**Figure S3.** Hydrogen bond (HB) propensity between unique donor-acceptor pairs for BglB-glucose complex (C0.1) over simulation time at 315 K ($T_{opt}$); yellow indicates HB presence, blue indicates absence. Right inset: time-dependent distance between glucose and Asn166 (−1 subsite), highlighting glucose expulsion from the catalytic pocket. Left inset: HB donor-acceptor pair while glucose resides in the active site (≤ 80 ns); later, glucose interacts with surface-exposed, non-catalytic residues. Across the three replicas glucose moved out within 100 ns. .....................................................................................8

**Figure S4.** Comparison of residue-wise glucose contact probabilities obtained using two distance cutoffs. Mean residue contact probability ($p_{ij}$) ± SEM across triplicate simulations is shown using a 3.8 Å hydrogen-bond cutoff (left; panels a–c) and a 7 Å contact cutoff (right; panels d–f). Rows correspond to glucose concentrations C1 (0.09 M), C2 (0.3 M), and C3 (1.0 M). The overall pattern of interacting residues and the resulting secondary binding sites (SBSs) were largely conserved between the two distance criteria, demonstrating that SBS identification was robust to the choice of contact cutoff. .......9

**Figure S5.** Residue-wise BglB-glucose interaction probability across concentrations. Panels (a–c) show the Heaviside-weighted consensus contact strength ($C_s^i$) calculated using a 3.8 Å contact cutoff for systems C1 (0.09 M), C2 (0.3 M), and C3 (1 M), respectively, whereas panels (d–f) show the corresponding analysis using a 7 Å contact cutoff. For each residue (i), ($C_s^i$) reflects the combined strength and reproducibility of residue–glucose interactions across three independent replicas (see Methods). ...............................................................................................................................10

**Figure S6.** Cumulative consensus contact strength ($C_{tot}$) for residues grouped into surface binding site (SBS) regions across systems C1-C3. For each residue i, the consensus contact ($\Sigma C_s^i$) strength was computed by summing the Heaviside-filtered fractional occupancies across three independent trajectories, such that only interactions with pij > 0.5 contributed. The cumulative SBS contact strength was then calculated as $C_{tot} = \Sigma C_s^i$ over all residues within each SBS. For comparison.......................11

Ctot calculated using $C_s^i$ derived from a 3.8 Å cutoff (a-c) and for a 7 Å cutoff (d-f) are shown. .........11

**Figure S7.** "Global" structural characteristics of wild-type (WT) and mutated SBS systems (mSBS1, mSBS2, mSBS4, and mSBS7) at 0 and 0.3 M glucose. (a, b) $C_\alpha$ root-mean-square fluctuation (RMSF); (c, d) backbone root-mean-square deviation (RMSD) over time; (e, f) solvent-accessible surface area (SASA) over time; (g, h) radius of gyration ($R_g$) over time; (i, j) secondary structure persistence over time. In each plot, lines represent mean values across three independent replica and shaded lines represent standard error of mean (SEM). ........ 12

**Figure S8.** "Global" and "Local" structural properties of WT and mutant (mSBS) systems. (a, b) solvent-accessible surface area (SASA); (c, d) secondary structure persistence ($S^p$); (e) Distance between catalytic residues Glu167 and Glu356 in WT and mSBS 1, mSBS2, mSBS4, and mSBS7. In each plot, lines represent mean values across three independent replica and shaded lines represent standard error of mean (SEM)........ 13

**Figure S9.** Residue-wise Protein- Glucose contact strength comparison between WT and mSBS variants. For each mutant (mSBS1, mSBS2, mSBS4, and mSBS7), the left panels (a-d) show the mean contact strength ($p_{ij}$) per residue, averaged over three independent replicas, with shaded (golden) regions representing the standard error of the mean (SEM). The right panels (e-h) display volcano plots of per-residue differences in contact strength ($\Delta p_{ij}$) versus statistical significance ($-\log_{10}$ p), where p-values are obtained from two-sided Welch's t-tests across replicas. Horizontal and vertical dashed lines denote the significance ($p < 0.05$) and change in ($|\Delta p_{ij}| > 0.1$; i.e., >10 %) cutoffs, respectively, and residues exceeding both thresholds are highlighted (deep teal). Mutated residues are further indicated with outlined circles........ 14

**Figure S10.** Residence times ($\tau$) were computed for water molecules within 7 Å of the entire protein surface ("Global") and (ii) the SBS region corresponding to each system. The first row shows $\tau$ at 0 M glucose, and the second row shows $\tau$ at 0.3 M glucose. (a, f) WT: global surface and SBS1, SBS2, SBS4, and SBS7 regions; (b, g) mSBS1: "Global" surface and mutated SBS1 region; (c, h) mSBS2: "Global" surface and mutated SBS2 region; (d, i) mSBS4: "Global" surface and mutated SBS4 region; (e, j) mSBS7: "Global" surface and mutated SBS7 region. These comparisons enable direct assessment of changes in water residence time both globally and at the mutated SBS regions relative to WT. ........ 15

**Figure S11.** Interaction energy decomposition of individual protein residues in WT C2 and the four mutant systems. Volcano plots showing the change in residue interaction energy ($\Delta$IE; kJ $mol^{-1}$) versus statistical significance ($-\log_{10} p$). The horizontal dashed line denotes the significance threshold ($p = 0.05$), while the vertical dashed lines indicate the interaction energy cutoff ($|\Delta IE| \geq 12.5$ kJ $mol^{-1}$), approximately equivalent to the strength of a hydrogen-bond interaction. Residues satisfying both criteria ($p < 0.05$ and $|\Delta IE| \geq 12.5$ kJ $mol^{-1}$) are highlighted. Panels correspond to (a) mSBS1, (b) mSBS2, (c) mSBS4, and (d) mSBS7; in each volcano plot, the residues mutated in the corresponding mutant are circled. (e–h) raw per-residue interaction energy contributions (mean ± SEM; $n$ = 3 independent simulations) for all residues satisfying both significance criteria, comparing WT C2 with (e) mSBS1, (f) mSBS2, (g) mSBS4, and (h) mSBS7, respectively. Mutated residues are marked with a golden star, whereas the remaining bars represent non-mutated residues exhibiting significant changes in interaction energy........................................................................................................................................ 16

**Figure S12.** PSN-based shortest paths connecting SBS regions to the catalytic core (active-site pocket, subsites −1 to +1). Comparison across WT at different glucose concentrations (C0-C3) shows that the shortest path lengths (and connecting nodes) are largely preserved. Mutant systems exhibited path lengths similar to WT under both conditions (with or without glucose). ......................................................... 17

**Figure S13.** Mutation-induced changes in community co-localization between the inner active-site pocket and surface SBS regions, expressed as the change in the number of residue pairs occupying the same interaction community relative to WT (mutant − WT). Diagonal elements represent local effects at the mutated SBS, whereas off-diagonal elements highlight long-range redistribution of communication pathways across the SBS network. Positive values indicate increased coupling between regions, whereas negative values indicate reduced community overlap and partial decoupling of communication pathways. The pronounced distal effects observed for mSBS2 and mSBS4 demonstrate that perturbations introduced at individual SBSs can propagate through pre-existing residue interaction networks and selectively reorganize communication with distant functional regions. ................................................ 18

**Figure S14.** (a) The Thermo Fisher protein ladder is shown in lane L1, and the eluted fraction of mSBS1 in lane L2 shows one additional protein band. (b) The Thermo Fisher protein ladder is shown in lane L4, and the eluted fraction of mSBS2 in lane L1 contains multiple protein bands, indicating lower sample purity. (c) The Thermo Fisher protein ladder is shown in lane L1, and the eluted fraction of mSBS7 in lane L2 shows a single predominant protein band, indicating successful purification. In each gel, a Thermo Fisher protein ladder was used, and the molecular weight marker is shown. .......................... 19

**Table S3.** Native BglB (WT) residues, their corresponding homologous residues identified through structure-based sequence alignment, and the final mutation used to generate each mob's variant. Structure-based sequence alignment of BglB (2O9P) with thermostable β-glucosidase homologs was performed to guide the design of secondary binding site (SBS) mutations. Homologs were selected from phylogenetically diverse bacterial and archaeal sources and including: 5DT5 from *Exiguobacterium antarcticum* B7[2] (BG1), 6RJK from *Agrobacterium tumefaciens* 5A[2] (BG2), 7E5J from *Thermoanaerobacterium saccharolyticum*[3] (TsaBgl) (BG3), 6IER from an uncultured bacterium[4] (BG4), 4PTV from *Halothermothrix orenii*[5] (BG5), 1GOW from *Saccharolobus solfataricus* P2[6] (BG6), 1UG6 from *Thermus thermophilus* (unpublished, 10.2210/pdb1ug6/pdb*)* (BG7), and 1VFF from *Pyrococcus horikoshii*[7] (BG8). The alignment was performed with the stride algorithm using VMD[8]. Residues that were 100% conserved across all homologs were retained without modification. For non-conserved positions within each SBS, the corresponding residues from homologs BG1-BG8 are listed in order (BG1/BG2/BG3/BG4/BG5/BG6/BG7/BG8). Positions labeled as "100% unconserved" indicate the absence of a consensus shared residue sequence among all homologs, in such cases, the residue from the most glucose-tolerant homolog is indicated in parentheses (e.g., GT-X). Substitution priority was given to *H. orenii* β-glucosidase[9] (4PTV), the homologs exhibiting the highest glucose tolerance (GT), followed by *A. tumefaciens* β-glucosidase (6RJK)[10]........................................ 23

**Table S4.** Residue-wise changes in glucose interaction energies for all mutated residues. The table reports the WT and mutant interaction energies (mean ± SEM; n = 3 independent simulations), the corresponding ΔIE (kJ $mol^{-1}$), and the normalized relative change in interaction energy. Positive relative ΔIE values indicate weaker interactions (loss), whereas negative values indicate stronger interactions (gain). Percentage values are provided only to illustrate the magnitude of energetic redistribution; for residues with interaction energies close to zero in the WT, percentage changes may be amplified by normalization. ........................................................................ 27

**Table S5.** Normalized Mutual Information (NMI), Adjusted Rand Index (ARI), and the fraction of residues retaining identical community membership following alignment to reference condition C0 (0 M glucose). Comparisons are shown both across glucose (GLC) concentrations (C0-C3) in the wild type (WT) and between WT (C0) and mutant structures (mSBS1, mSBS2, mSBS4 and mSBS7). Based on alignment of consensus community across three replicas…………………………………………..... 28

**Table S6.** Summary of system preparation for BglB simulations, showing glucose concentration (M), number of glucose molecules ($N_{Glucose}$), and number of water molecules ($N_{Water}$)................................ 29

**Figure S1.** Changes in relative activity of BglB with increasing concentrations of exogenous glucose (0-1M).

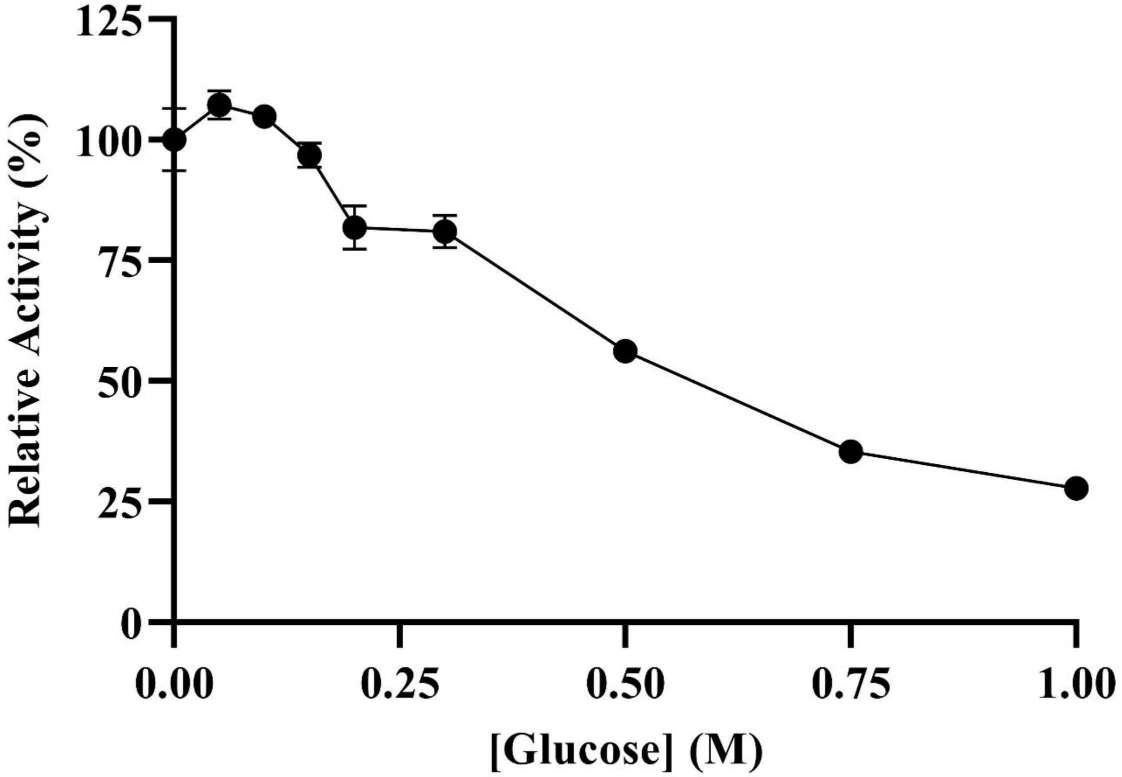

**Figure S2.** "Global" Structural characteristics of BglB in different glucose concentrations. Backbone RMSD timeline, C-alpha RMSF (highlighted gatekeeper residues), secondary structure persistence and SASA timeline for BglB. In each plot, lines represent mean values across three independent replica and shaded lines represent standard error of mean (SEM).

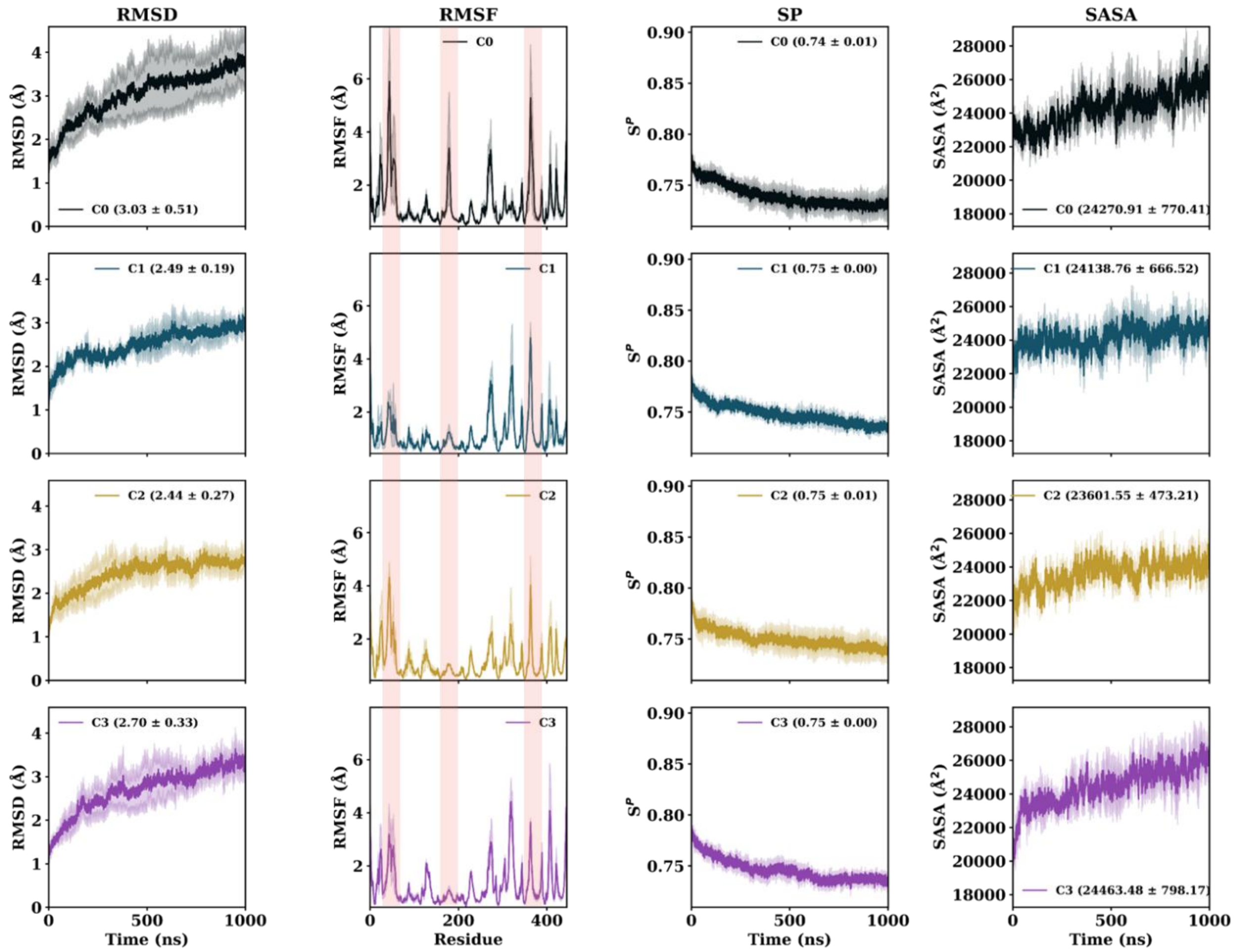

**Figure S3.** Hydrogen bond (HB) propensity between unique donor-acceptor pairs for BglB-glucose complex (C0.1) over simulation time at 315 K ($T_{opt}$); yellow indicates HB presence, blue indicates absence. Right inset: time-dependent distance between glucose and Asn166 (−1 subsite), highlighting glucose expulsion from the catalytic pocket. Left inset: HB donor-acceptor pair while glucose resides in the active site (≤ 80 ns); later, glucose interacts with surface-exposed, non-catalytic residues. Across the three replicas glucose moved out within 100 ns.

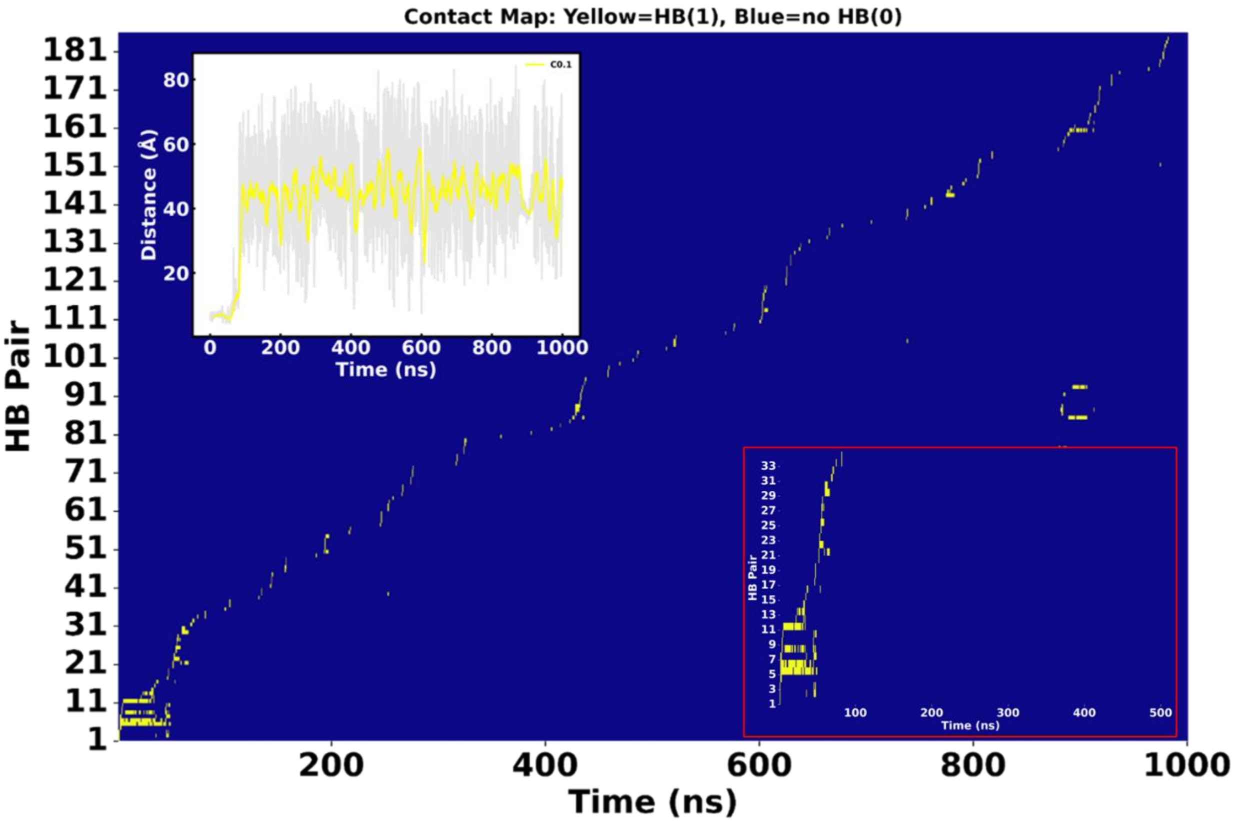

**Figure S4.** Comparison of residue-wise glucose contact probabilities obtained using two distance cutoffs. Mean residue contact probability ($p_{ij}$) ± SEM across triplicate simulations is shown using a 3.8 Å hydrogen-bond cutoff (left; panels a–c) and a 7 Å contact cutoff (right; panels d–f). Rows correspond to glucose concentrations C1 (0.09 M), C2 (0.3 M), and C3 (1.0 M). The overall pattern of interacting residues and the resulting secondary binding sites (SBSs) were largely conserved between the two distance criteria, demonstrating that SBS identification was robust to the choice of contact cutoff.

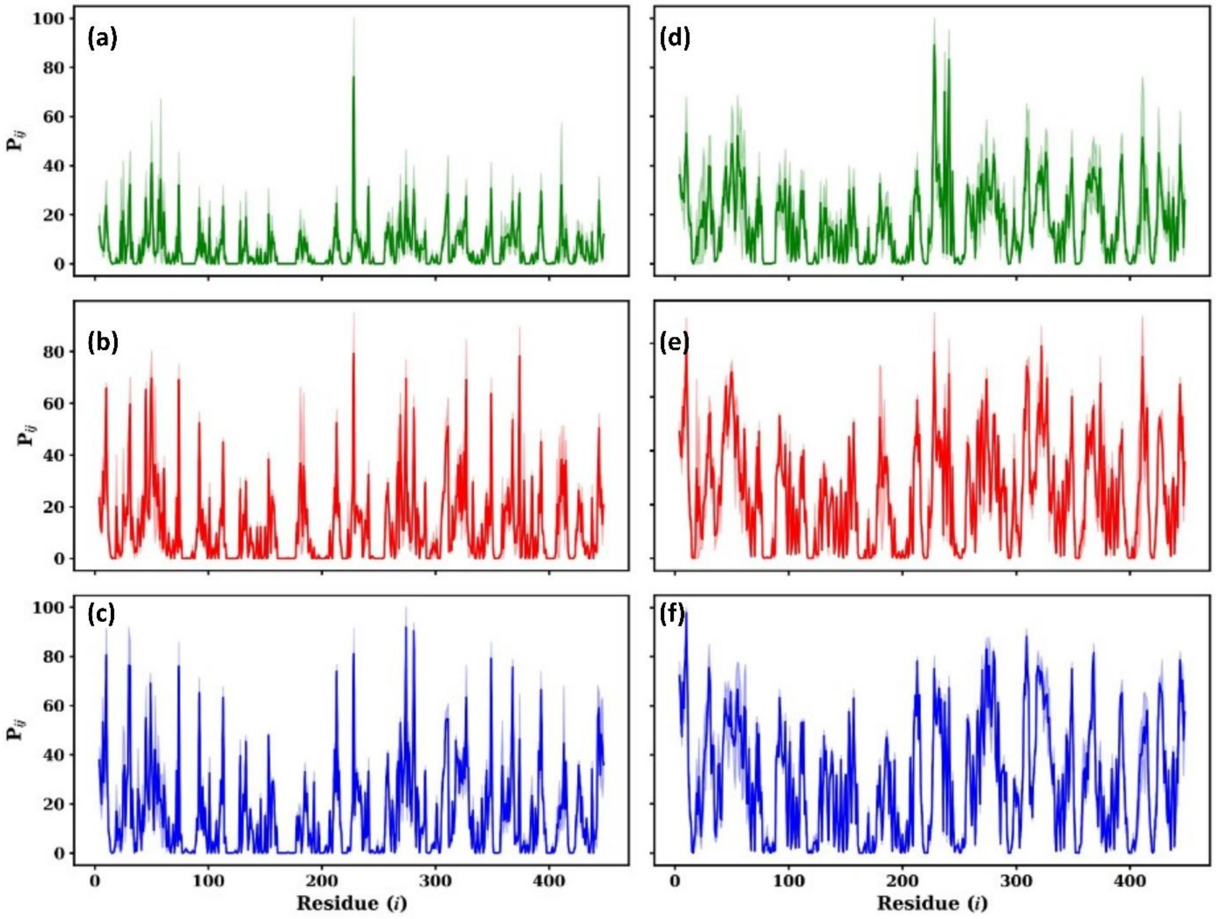

**Figure S5.** Residue-wise BglB-glucose interaction probability across concentrations. Panels (a–c) show the Heaviside-weighted consensus contact strength ($C_S^i$) calculated using a 3.8 Å contact cutoff for systems C1 (0.09 M), C2 (0.3 M), and C3 (1 M), respectively, whereas panels (d–f) show the corresponding analysis using a 7 Å contact cutoff. For each residue (i), ($C_S^i$) reflects the combined strength and reproducibility of residue–glucose interactions across three independent replicas (see Methods).

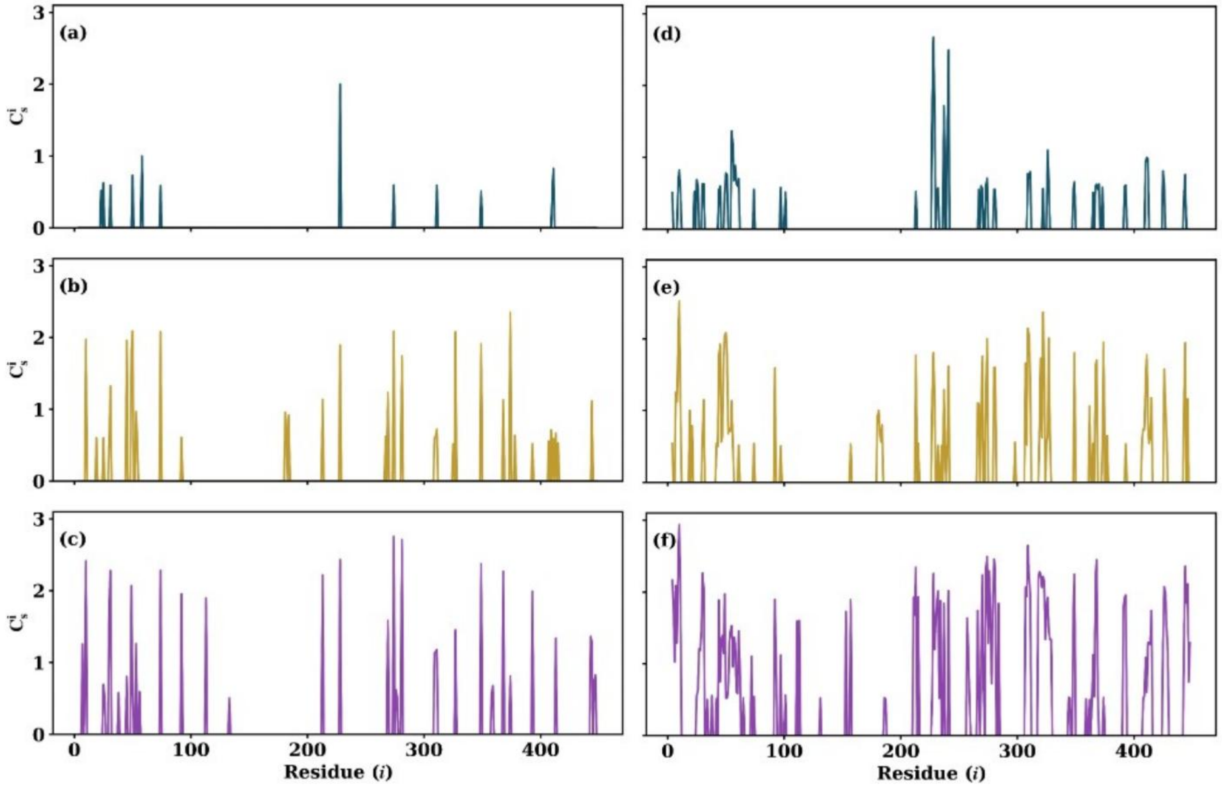

**Figure S6.** Cumulative consensus contact strength ($C_{\text{tot}}$) for residues grouped into surface binding site (SBS) regions across systems C1-C3. For each residue i, the consensus contact ($\Sigma C_s^i$) strength was computed by summing the Heaviside-filtered fractional occupancies across three independent trajectories, such that only interactions with $p_{ij}$ > 0.5 contributed. The cumulative SBS contact strength was then calculated as $C_{\text{tot}} = \Sigma C_s^i$ over all residues within each SBS. For comparison $C_{\text{tot}}$ calculated using $C_s^i$ derived from a 3.8 Å cutoff (a-c) and for a 7 Å cutoff (d-f) are shown.

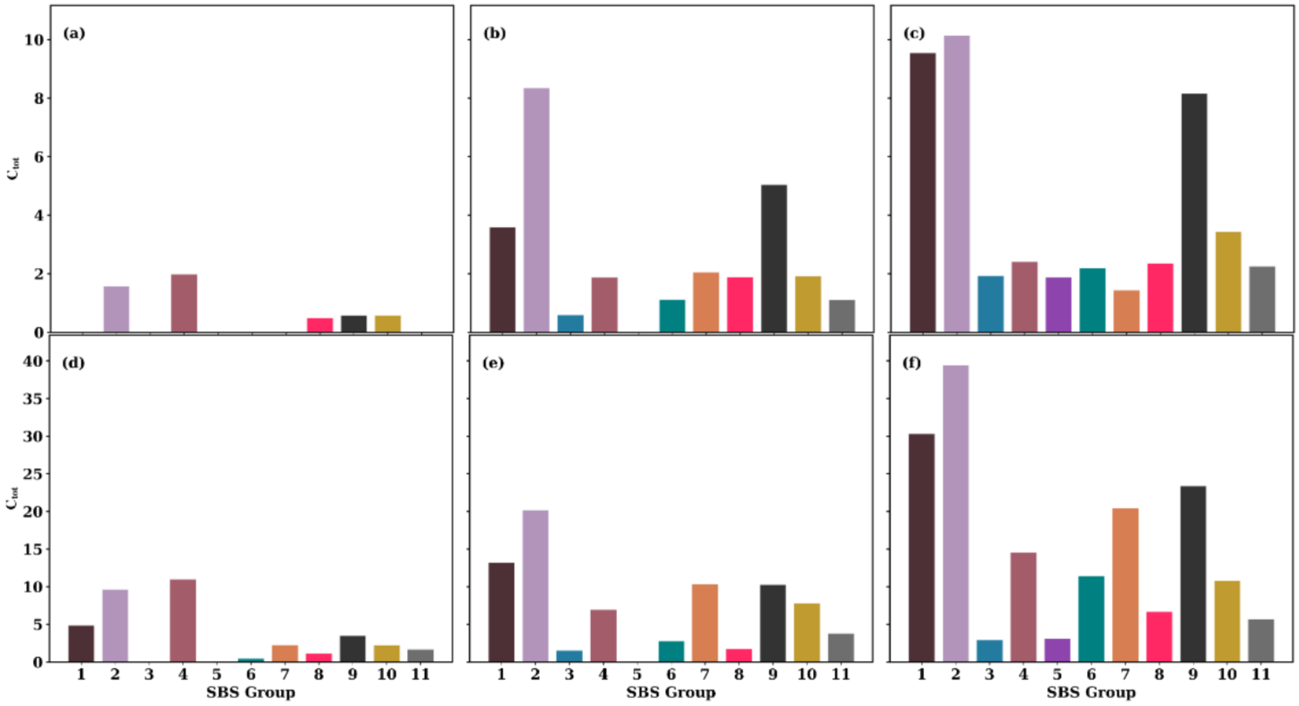

**Figure S7.** "Global" structural characteristics of wild-type (WT) and mutated SBS systems (mSBS1, mSBS2, mSBS4, and mSBS7) at 0 and 0.3 M glucose. (a, b) $C_\alpha$ root-mean-square fluctuation (RMSF); (c, d) backbone root-mean-square deviation (RMSD) over time; (e, f) solvent-accessible surface area (SASA) over time; (g, h) radius of gyration ($R_g$) over time; (i, j) secondary structure persistence over time. In each plot, lines represent mean values across three independent replica and shaded lines represent standard error of mean (SEM).

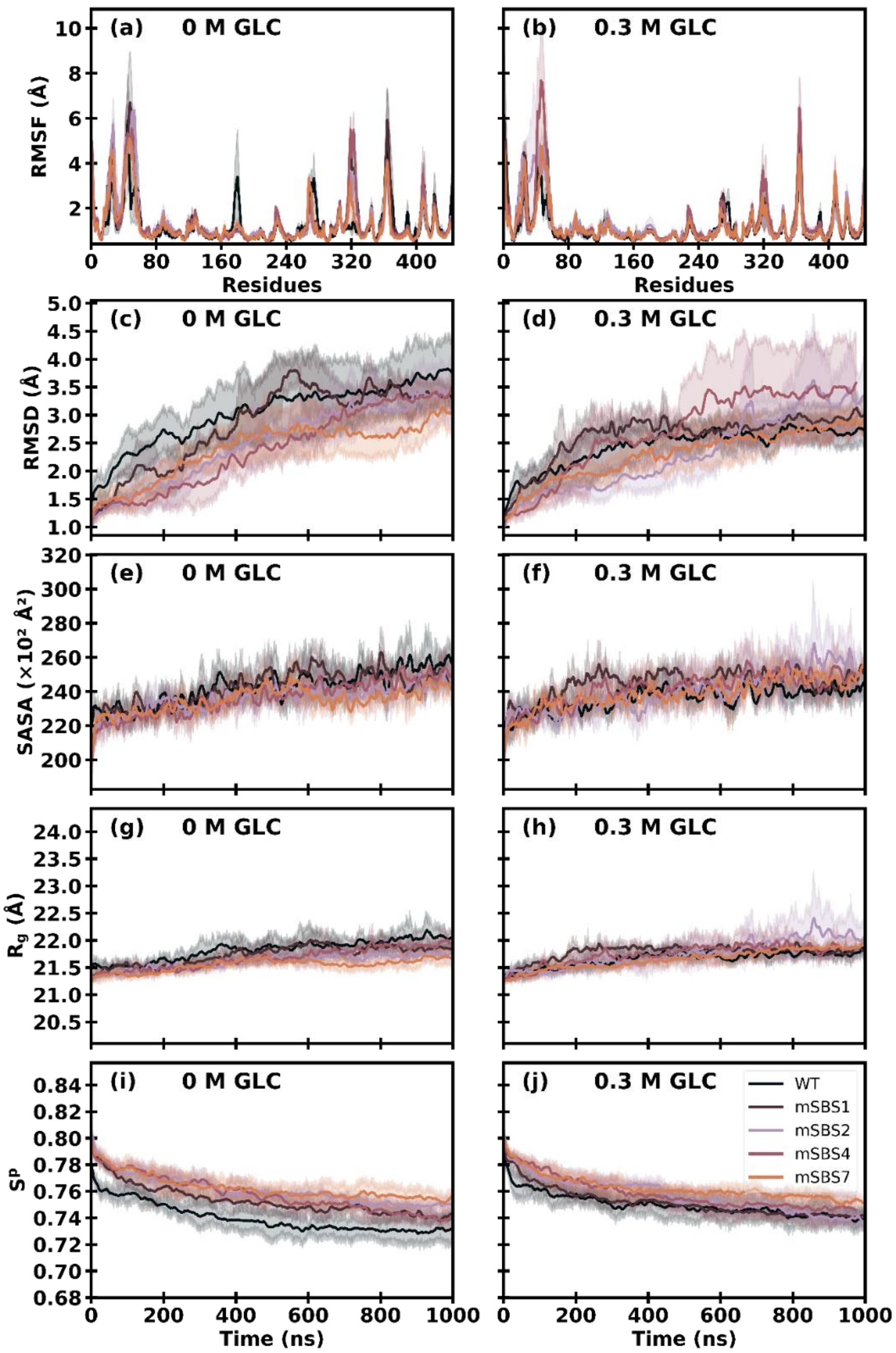

**Figure S8.** "Global" and "Local" structural properties of WT and mutant (mSBS) systems. (a, b) solvent-accessible surface area (SASA); (c, d) secondary structure persistence ($S^p$); (e) Distance between catalytic residues Glu167 and Glu356 in WT and mSBS 1, mSBS2, mSBS4, and mSBS7. In each plot, lines represent mean values across three independent replica and shaded lines represent standard error of mean (SEM).

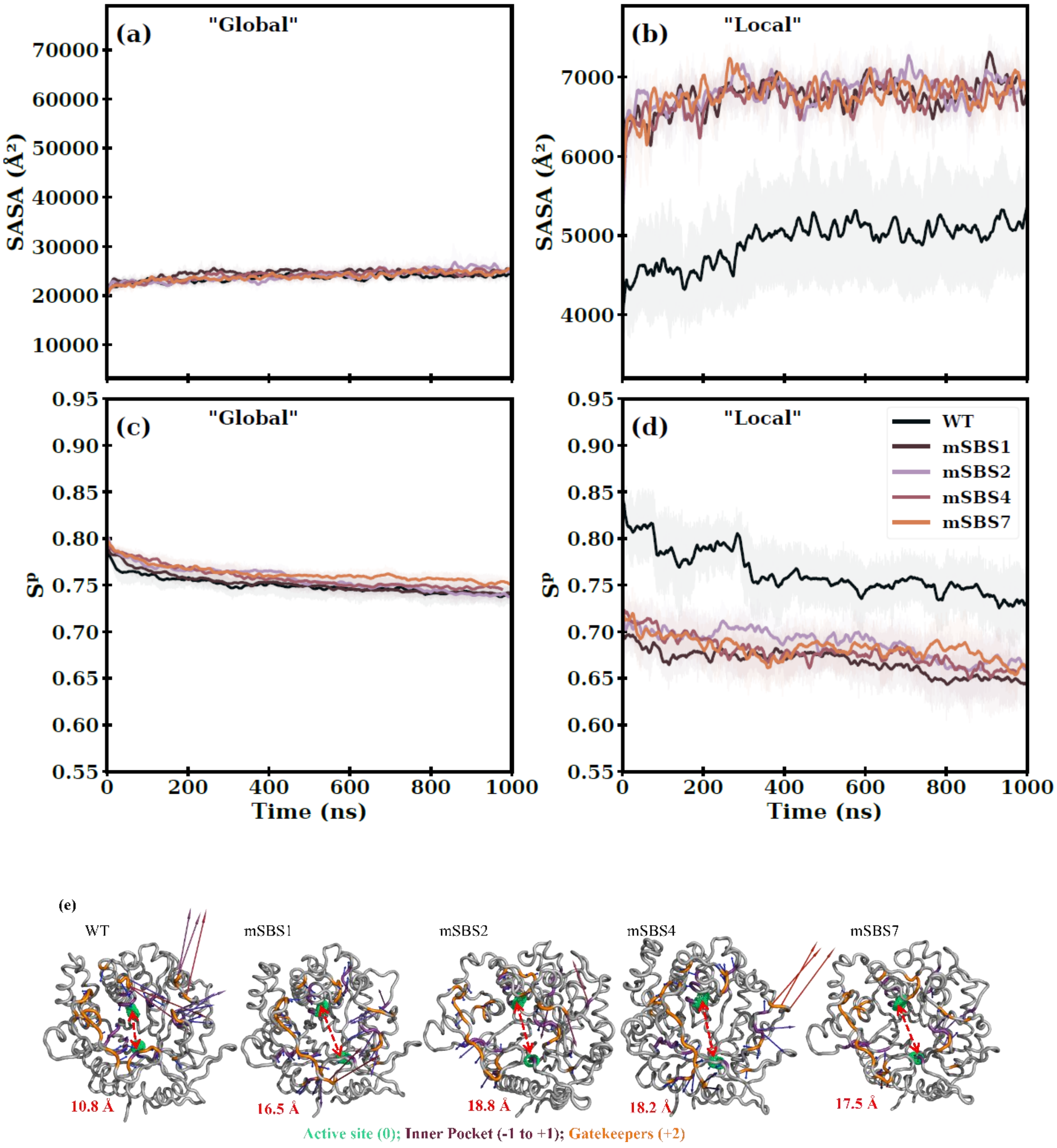

**Figure S9.** Residue-wise Protein- Glucose contact strength comparison between WT and mSBS variants. For each mutant (mSBS1, mSBS2, mSBS4, and mSBS7), the left panels (a-d) show the mean contact strength ($p_{ij}$) per residue, averaged over three independent replicas, with shaded (golden) regions representing the standard error of the mean (SEM). The right panels (e-h) display volcano plots of per-residue differences in contact strength ($\Delta p_{ij} = P_{ij}^{\mathrm{mSBS}} - P_{ij}^{\mathrm{WT}}$) versus statistical significance ($-\log_{10}$ p), where p-values are obtained from two-sided Welch's t-tests across replicas. Horizontal and vertical dashed lines denote the significance ($p < 0.05$) and change in ($|\Delta p_{ij}| > 0.1$; i.e., >10 %) cutoffs, respectively, and residues exceeding both thresholds are highlighted (deep teal). Mutated residues are further indicated with outlined circles.

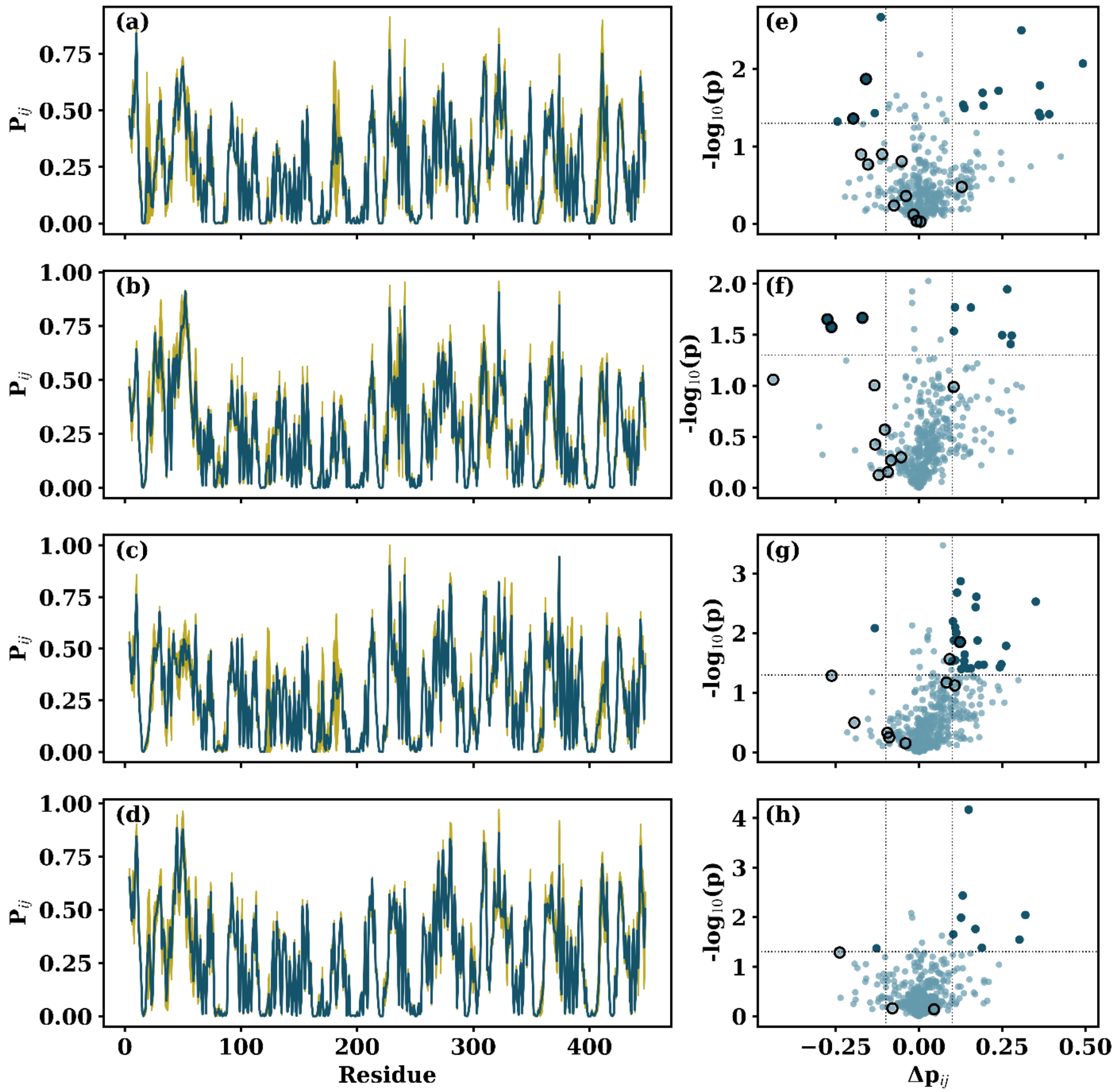

**Figure S10.** Residence times (τ) were computed for water molecules within 7 Å of the entire protein surface ("Global") and (ii) the SBS region corresponding to each system. The first row shows τ at 0 M glucose, and the second row shows τ at 0.3 M glucose. (a, f) WT: global surface and SBS1, SBS2, SBS4, and SBS7 regions; (b, g) mSBS1: "Global" surface and mutated SBS1 region; (c, h) mSBS2: "Global" surface and mutated SBS2 region; (d, i) mSBS4: "Global" surface and mutated SBS4 region; (e, j) mSBS7: "Global" surface and mutated SBS7 region. These comparisons enable direct assessment of changes in water residence time both globally and at the mutated SBS regions relative to WT.

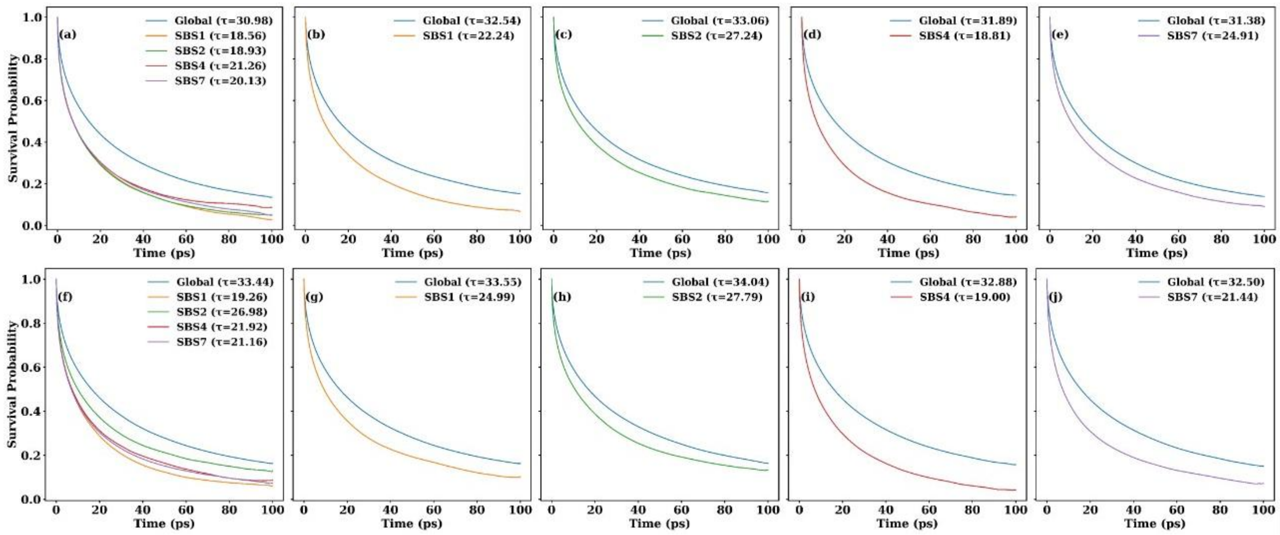

**Figure S11.** Per-residue interaction energy decomposition of individual protein residues in WT C2 and the four mutant systems. Volcano plots showing the change in residue interaction energy ($\Delta IE = IE_{\text{mutant}} - IE_{\text{WT}}$; kJ $mol^{-1}$) versus statistical significance ($-\log_{10} p$). The horizontal dashed line denotes the significance threshold ($p = 0.05$), while the vertical dashed lines indicate the interaction energy cutoff ($|\Delta IE| \geq 12.5$ kJ $mol^{-1}$), approximately equivalent to the strength of a hydrogen-bond interaction. Residues satisfying both criteria ($p < 0.05$ and $|\Delta IE| \geq 12.5$ kJ $mol^{-1}$) are highlighted. Panels correspond to (a) mSBS1, (b) mSBS2, (c) mSBS4, and (d) mSBS7; in each volcano plot, the residues mutated in the corresponding mutant are circled. (e–h) raw per-residue interaction energy contributions (mean ± SEM; $n = 3$ independent simulations) for all residues satisfying both significance criteria, comparing WT C2 with (e) mSBS1, (f) mSBS2, (g) mSBS4, and (h) mSBS7, respectively. Mutated residues are marked with a golden star, whereas the remaining bars represent non-mutated residues exhibiting significant changes in interaction energy.

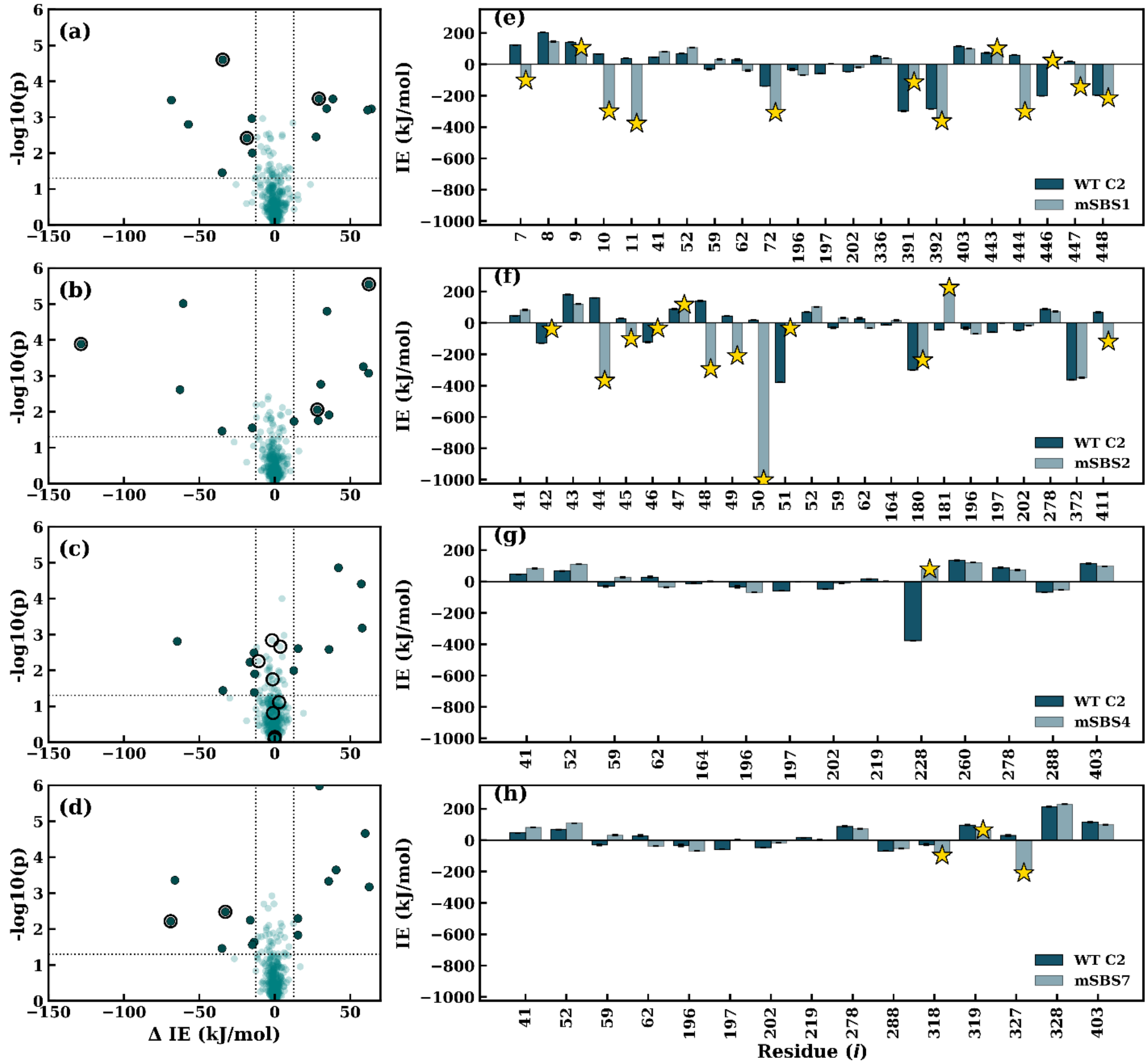

**Figure S12.** PSN-based shortest paths connecting SBS regions to the catalytic core (active-site pocket, subsites −1 to +1). Comparison across WT at different glucose concentrations (C0-C3) shows that the shortest path lengths (and connecting nodes) are largely preserved. Mutant systems exhibited path lengths similar to WT under both conditions (with or without glucose).

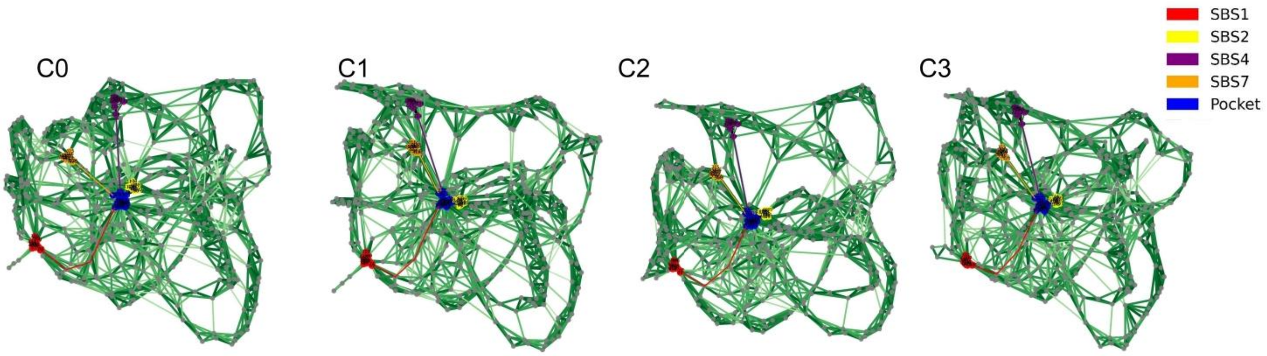

**Figure S13.** Mutation-induced changes in community co-localization between the inner active-site pocket and surface SBS regions, expressed as the change in the number of residue pairs occupying the same interaction community relative to WT (Δ co-localization = mutant − WT). Diagonal elements represent local effects at the mutated SBS, whereas off-diagonal elements highlight long-range redistribution of communication pathways across the SBS network. Positive values indicate increased coupling between regions, whereas negative values indicate reduced community overlap and partial decoupling of communication pathways. The pronounced distal effects observed for mSBS2 and mSBS4 demonstrate that perturbations introduced at individual SBSs can propagate through pre-existing residue interaction networks and selectively reorganize communication with distant functional regions.

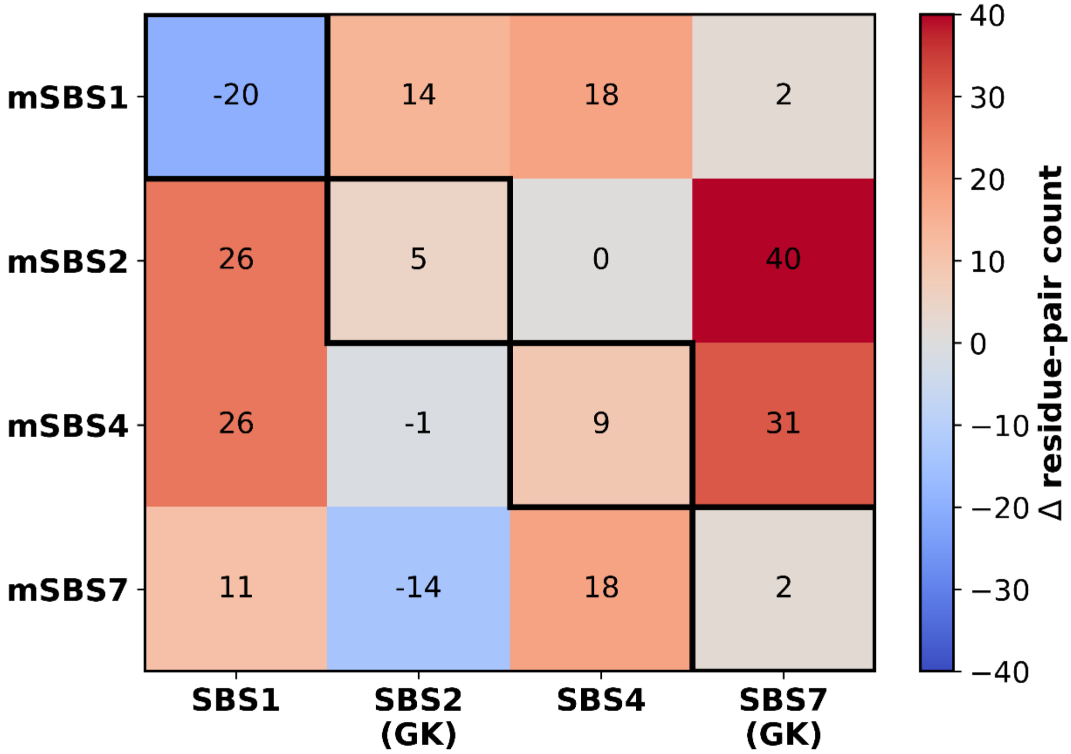

**Figure S14.** (a) The Thermo Fisher protein ladder is shown in lane L1, and the eluted fraction of mSBS1 in lane L2 shows one additional protein band. (b) The Thermo Fisher protein ladder is shown in lane L4, and the eluted fraction of mSBS2 in lane L1 contains multiple protein bands, indicating lower sample purity. (c) The Thermo Fisher protein ladder is shown in lane L1, and the eluted fraction of mSBS7 in lane L2 shows a single predominant protein band, indicating successful purification. In each gel, a Thermo Fisher protein ladder was used, and the molecular weight marker is shown.

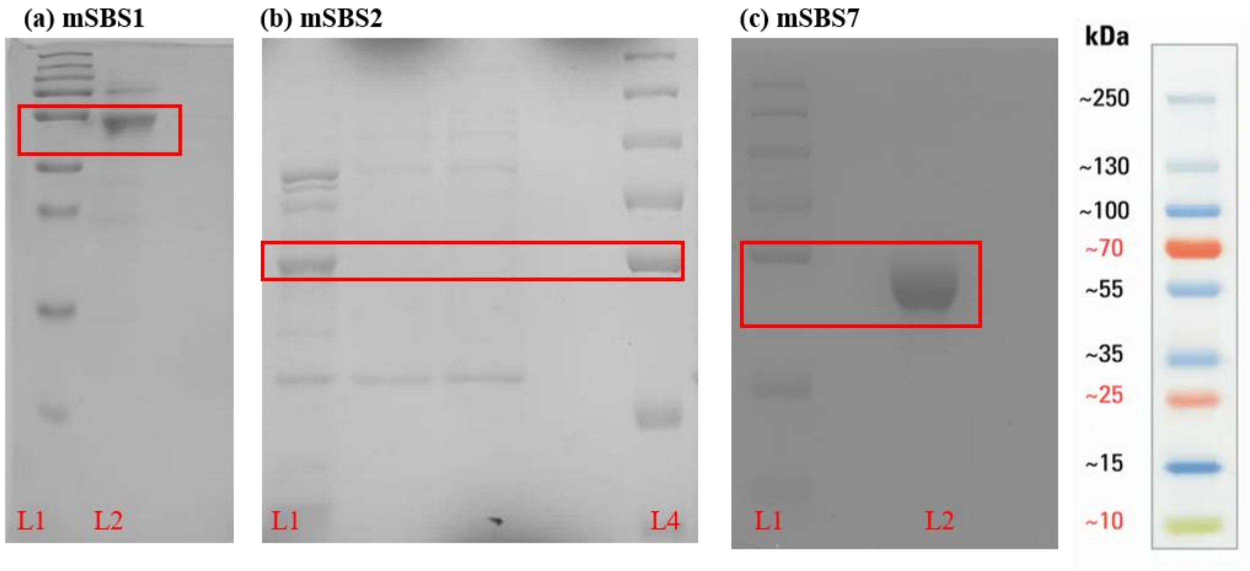

**Figure S15.** Circular dichroism (CD) spectra of WT and mutant systems (mSBS1 and mSBS7). The corresponding table reports the percentage secondary structure (% SS) content estimated using the BeStSel server[1].

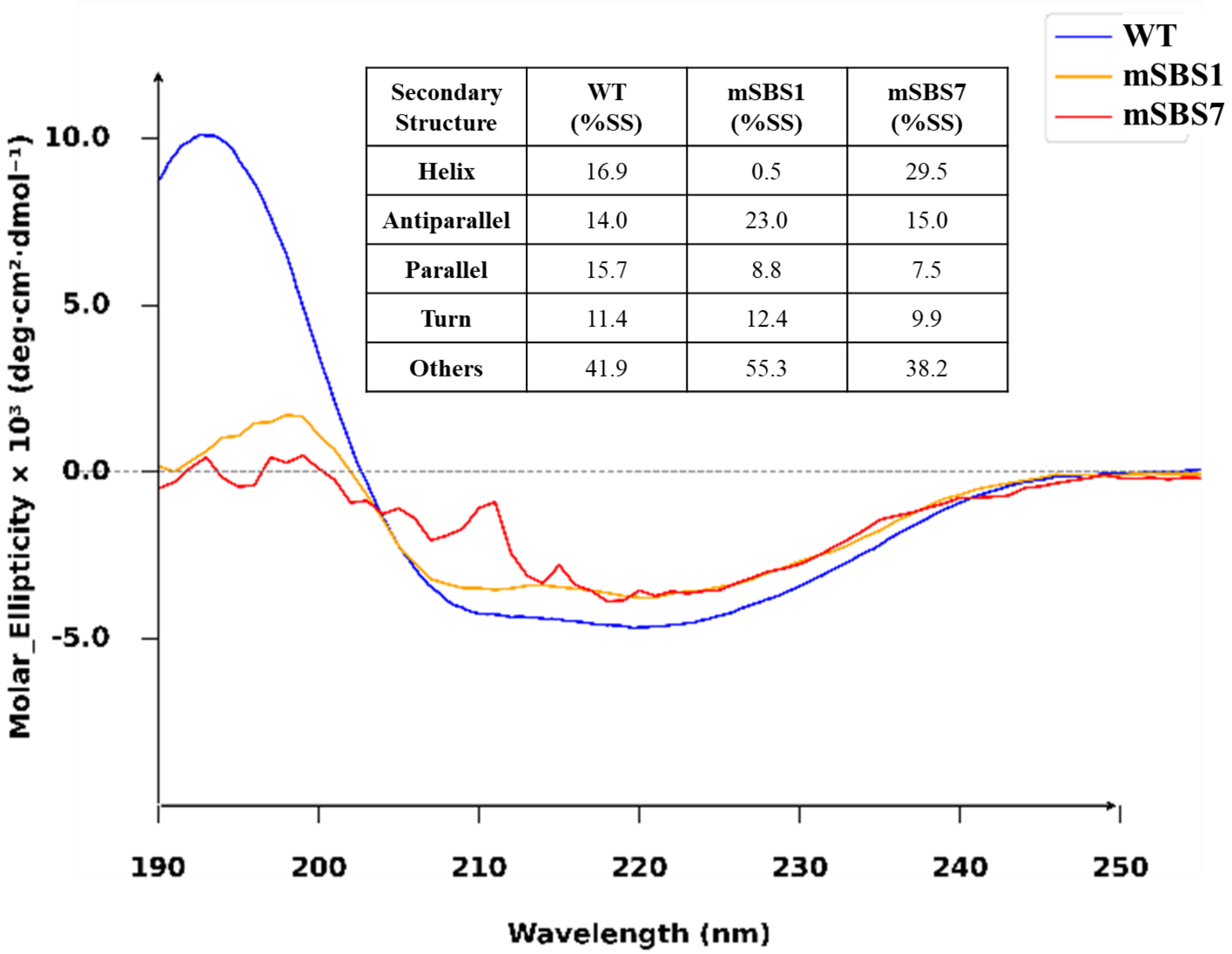

| Secondary Structure | WT (%SS) | mSBS1 (%SS) | mSBS7 (%SS) |
|---|---|---|---|
| Helix | 16.9 | 0.5 | 29.5 |
| Antiparallel | 14.0 | 23.0 | 15.0 |
| Parallel | 15.7 | 8.8 | 7.5 |
| Turn | 11.4 | 12.4 | 9.9 |
| Others | 41.9 | 55.3 | 38.2 |

**Table S1**. $K_m$ and $V_{max}$ of WT and mSBS7 mutant from non-linear regression analysis (Michaelis-Menten Kinetics) with exogenous glucose (0-0.5 M) on *p*NPGlc.

| **WT** | | | | | | | |
|---|---|---|---|---|---|---|---|
| **Glucose (M)** | 0 | 0.050 | 0.10 | 0.15 | 0.20 | 0.30 | 0.50 |
| **$K_m$ (mM)** | 10.16 | 13.23 | 25.16 | 33.57 | 42.08 | 42.96 | 64.71 |
| **$V_{max}$ (mM $min^{-1}$)** | 0.22 | 0.24 | 0.26 | 0.26 | 0.24 | 0.23 | 0.20 |
| **mSBS7** | | | | | | | |
| **Glucose (M)** | 0 | 0.050 | 0.10 | 0.15 | 0.20 | 0.30 | 0.50 |
| **$K_m$ (mM)** | 6.39 | 8.16 | 14.09 | 19.09 | 22.49 | 22.15 | 46.23 |
| **$V_{max}$ (mM $min^{-1}$)** | 0.14 | 0.16 | 0.16 | 0.16 | 0.15 | 0.14 | 0.13 |

**Table S2.** Secondary binding sites (SBSs) and participating residues. Residues identified from $C_s^i$ and mapped onto the protein surface, forming distinct surface clusters, are listed as secondary binding sites (SBSs).

| SBS Index | Number of Residues involved | Residue IDs |
|---|---|---|
| 1 | 16 | ASN4, THR5, PHE6, ILE7, PHE8, PRO9, ALA10, THR11, GLU391, GLU392, GLY393, MET443, ALA444, LYS445, ASN446, PHE448 |
| 2 | 27 | THR27, ASP28, GLU29, GLY30, GLY31, GLY53, ASP54, VAL55, ALA56, CYS57, ASP58, HIS61, GLU97, PRO44, GLY45, LYS46, VAL47, ILE48, GLY49, GLY413, TYR414, SER415, TYR425, GLU426, THR427, GLN428, GLU429 |
| 3 | 2 | GLY92, ILE93 |
| 4 | 9 | ASP228, ALA229, ALA230, GLU232, PRO234, SER231, VAL237, ALA240, ILE241 |
| 5 | 2 | LEU111, GLY113 |
| 6 | 6 | ASP153, GLU157, GLU211, LYS212, GLY213, THR215 |
| 7 | 11 | HIS318, MET319, GLU320, GLU321, PRO322, VAL323, THR324, MET326, GLY327, TRP328, GLU329 |
| 8 | 4 | ASN257, GLY258, LYS348, GLY349 |
| 9 | 12 | GLU266, GLY269, THR270, ASN273, GLY274, LEU275, ASP276, PHE277, GLN279, PRO280, GLY281, GLU284 |
| 10 | 5 | ASN307, ASP308, ALA309, SER310, LEU311 |
| 11 | 3 | LEU365, ASN367, GLY368 |

**Table S3.** Native BglB (WT) residues, their corresponding homologous residues identified through structure-based sequence alignment, and the final mutation used to generate each mob's variant. Structure-based sequence alignment of BglB (2O9P) with thermostable β-glucosidase homologs was performed to guide the design of secondary binding site (SBS) mutations. Homologs were selected from phylogenetically diverse bacterial and archaeal sources and including: 5DT5 from *Exiguobacterium antarcticum* B7[2] (BG1), 6RJK from *Agrobacterium tumefaciens* 5A[2] (BG2), 7E5J from *Thermoanaerobacterium saccharolyticum*[3] (TsaBgl) (BG3), 6IER from an uncultured bacterium[4] (BG4), 4PTV from *Halothermothrix orenii*[5] (BG5), 1GOW from *Saccharolobus solfataricus* P2[6] (BG6), 1UG6 from *Thermus thermophilus* (unpublished, 10.2210/pdb1ug6/pdb*)* (BG7), and 1VFF from *Pyrococcus horikoshii*[7] (BG8). The alignment was performed with the stride algorithm using VMD[8]. Residues that were 100% conserved across all homologs were retained without modification. For non-conserved positions within each SBS, the corresponding residues from homologs BG1-BG8 are listed in order (BG1/BG2/BG3/BG4/BG5/BG6/BG7/BG8). Positions labeled as "100% unconserved" indicate the absence of a consensus shared residue sequence among all homologs, in such cases, the residue from the most glucose-tolerant homolog is indicated in parentheses (e.g., GT-X). Substitution priority was given to *H. orenii* β-glucosidase[9] (4PTV), the homologs exhibiting the highest glucose tolerance (GT), followed by *A. tumefaciens* β-glucosidase (6RJK)[10].

| **mSBS1** | | |
|---|---|---|
| **SBS1** | **Residues in homologous BG (BG1/BG2/BG3/BG4/BG5/BG6/BG7/BG8)** | **mutated based on 4PTV/6RJK (BG2/BG5) Mutant SBS1** |
| I7 | 100% unconserved (GT-K) | I7K |
| P9 | S/A (GT-S) | A9S |
| A10 | K/E/P/G/N | T10E |
| Q72 | D/E/S/K | Q72E |
| E391 | N/K/D/R | E391K |

| | | |
|---|---|---|
| E392 | D/S | E392D |
| M443 | I/A | M443I |
| A444 | S/E/A/Q/R/T | A444E |
| K445 | T/R/G/S/A/N | K445K |
| N446 | R/G/A/H/F/K | N446G |
| G447 | 100% unconserved (GT-Q) | G447Q |
| F448 | 100% unconserved (GT-V) | F448V |
| **mSBS2** | | |
| **SBS2** | **Residues in homologous BG**<br>**(BG1/BG2/BG3/BG4/BG5/BG6/BG7/BG8** | **mutated based on 4PTV/6RJK**<br>**Mutant SBS2** |
| Q42 | R/H//D/N//H/H | Q42H |
| P44 | D///D//G/Q/A | P44D |
| G45 | K/T | G45K |
| K46 | I/A/H/L | K46H |

| | | |
|---|---|---|
| V47 | I/P/F | V47I |
| I48 | Y/E/R/F/E/S | I48E |
| G49 | 100% unconserved (GT-N) | G49N |
| G50 | S/G/K/R | G50R |
| D51 | H/S/R/E/D | D51H |
| E180 | N/E/Q/I/Y/S | E180N |
| H181 | W/V/F | H181W |
| A411 | S/K/F | A411K |
| **mSBS4** | | |
| **SBS4** | **Residues in homologous BG (BG1/BG2/BG3/BG4/BG5/BG6/BG7/BG8** | **mutated based on 4PTV/6RJK Mutant SBS4** |
| V227 | A/K/E/F | V227A |
| D228 | F/Y/I/L/R/Q | D228Y |
| A229 | P | A229P |

| | | |
|---|---|---|
| A230 | G/K/A | A230G |
| 231 | G/T/S/E | S231G |
| E232 | D/N/P | E232D |
| P234 | 100 % unconserved (GT-E) | P234E |
| V237 | K/P/Q/M/R/A | V237M |
| I241 | Q/S/D/N/E/D | I241S |
| **mSBS7** | | |
| **SBS7** | **Residues in homologous BG (BG1/BG2/BG3/BG4/BG5/BG6/BG7/BG8** | **mutated based on 4PTV/6RJK Mutant SBS7** |
| H318 | D/K/P/Y/L/N/G | H318K |
| M319 | P/M/P/S/A/P/A | M319A |
| G327 | N | G327N |

**Table S4.** Residue-wise changes in glucose interaction energies for all mutated residues. The table reports the WT and mutant interaction energies (mean ± SEM; n = 3 independent simulations), the corresponding ΔIE (kJ mol$^{-1}$), and the normalized relative change in interaction energy. Positive relative ΔIE values indicate weaker interactions (loss), whereas negative values indicate stronger interactions (gain). Percentage values are provided only to illustrate the magnitude of energetic redistribution; for residues with interaction energies close to zero in the WT, percentage changes may be amplified by normalization.

| **Mutant** | **Residue** | **WT IE (Mean ± SEM)** | **Mutant IE (Mean ± SEM)** | **ΔIE (kJ mol$^{-1}$)** | **Relative ΔIE (%)** | **Interpret\ation** |
|---|---|---|---|---|---|---|
| **mSBS1** | 7 | −0.49 ± 0.06 | 0.01 ± 0.07 | 0.50 | 102.7 | Loss |
| | 9 | −0.39 ± 0.03 | −0.16 ± 0.03 | 0.23 | 58.6 | Loss |
| | 10 | −0.39 ± 0.02 | 0.12 ± 0.02 | 0.51 | 130.2 | Loss |
| | 11 | −0.26 ± 0.02 | −0.01 ± 0.04 | 0.25 | 95.6 | Loss |
| | 72 | −0.31 ± 0.03 | 0.13 ± 0.02 | 0.44 | 141.4 | Loss |
| | 391 | 0.21 ± 0.01 | 0.11 ± 0.01 | −0.09 | −44.7 | Gain |
| | 392 | 0.25 ± 0.01 | 0.27 ± 0.01 | 0.02 | 9.7 | Loss |
| | 443 | −0.49 ± 0.07 | −0.34 ± 0.05 | 0.15 | 31.1 | Loss |
| | 444 | −0.23 ± 0.01 | 0.15 ± 0.01 | 0.38 | 165.0 | Loss |
| | 446 | −0.37 ± 0.04 | −0.15 ± 0.04 | 0.23 | 61.2 | Loss |
| | 447 | −0.06 ± 0.03 | −0.22 ± 0.05 | −0.16 | −287.9 | Gain |
| | 448 | −0.10 ± 0.04 | 0.15 ± 0.10 | 0.25 | 239.0 | Loss |
| **mSBS2** | 42 | −0.25 ± 0.03 | −0.16 ± 0.02 | 0.09 | 35.9 | Loss |
| | 44 | −0.39 ± 0.02 | 0.23 ± 0.03 | 0.62 | 157.7 | Loss |
| | 45 | −0.15 ± 0.01 | 0.10 ± 0.04 | 0.25 | 164.9 | Loss |
| | 46 | 0.08 ± 0.02 | −0.11 ± 0.05 | −0.19 | −239.6 | Gain |
| | 47 | −0.35 ± 0.06 | −0.27 ± 0.08 | 0.07 | 21.4 | Loss |
| | 48 | −0.46 ± 0.03 | 0.22 ± 0.01 | 0.68 | 146.5 | Loss |
| | 49 | −0.15 ± 0.03 | −0.20 ± 0.03 | −0.05 | −30.4 | Gain |
| | 50 | −0.19 ± 0.01 | −0.26 ± 0.04 | −0.06 | −32.8 | Gain |
| | 51 | 0.20 ± 0.03 | −0.23 ± 0.00 | −0.43 | −212.9 | Gain |
| | 180 | 0.29 ± 0.05 | −0.20 ± 0.03 | −0.50 | −169.6 | Gain |
| | 181 | −0.25 ± 0.21 | −0.65 ± 0.53 | −0.40 | −161.2 | Gain |
| | 411 | −0.36 ± 0.07 | 0.04 ± 0.13 | 0.40 | 111.7 | Loss |
| **mSBS4** | 227 | −0.44 ± 0.01 | −0.16 ± 0.05 | 0.28 | 64.4 | Loss |
| | 228 | 0.13 ± 0.02 | −0.51 ± 0.03 | −0.64 | −485.0 | Gain |
| | 229 | −0.23 ± 0.05 | −0.18 ± 0.01 | 0.04 | 19.4 | Loss |
| | 230 | −0.11 ± 0.00 | −0.14 ± 0.00 | −0.03 | −24.7 | Gain |
| | 231 | −0.09 ± 0.01 | −0.10 ± 0.00 | −0.02 | −19.8 | Gain |
| | 232 | 0.24 ± 0.01 | 0.26 ± 0.00 | 0.02 | 9.9 | Loss |
| | 234 | −0.40 ± 0.01 | −0.39 ± 0.01 | 0.00 | 1.1 | Loss |
| | 237 | −0.44 ± 0.07 | −0.37 ± 0.00 | 0.08 | 17.5 | Loss |
| | 241 | −0.63 ± 0.09 | −0.54 ± 0.06 | 0.10 | 15.2 | Loss |
| **mSBS7** | 318 | −0.17 ± 0.03 | 0.07 ± 0.02 | 0.23 | 138.8 | Loss |
| | 319 | −0.57 ± 0.09 | −0.23 ± 0.06 | 0.34 | 60.0 | Loss |
| | 327 | −0.17 ± 0.01 | −0.21 ± 0.03 | −0.04 | −24.3 | Gain |

**Table S5.** Normalized Mutual Information (NMI), Adjusted Rand Index (ARI), and the fraction of residues retaining identical community membership following alignment to reference condition C0 (0 M glucose). Comparisons are shown both across glucose (GLC) concentrations (C0-C3) in the wild type (WT) and between WT (C0) and mutant structures (mSBS1, mSBS2, mSBS4 and mSBS7). Based on alignment of consensus community across three replicas.

| | **NMI** | **ARI** | **VI** |
|---|---|---|---|
| **C0, C1** | 0.77 | 0.63 | 0.90 |
| **C0, C2** | 0.87 | 0.79 | 0.00 |
| **C0, C3** | 0.83 | 0.75 | 1.02 |
| **C0, mSBS1** | 0.70 | 0.55 | 1.34 |
| **C0, mSBS2** | 0.74 | 0.60 | 0.76 |
| **C0, mSBS4** | 0.70 | 0.51 | 0.95 |
| **C0, mSBS7** | 0.72 | 0.66 | 0.90 |

**Table S6.** Summary of system preparation for BglB simulations, showing glucose concentration (M), number of glucose molecules ($N_{Glucose}$), and number of water molecules ($N_{Water}$).

| System | Glucose (M) | $N_{Glucose}$ | $N_{Water}$ | % Relative Specific Activity |
|---|---|---|---|---|
| C0 | 0.00 | 0 | 20689 | 100 |
| C0.1 | 0.00 | 1 | 20616 | - |
| C1 | 0.09 | 39 | 19883 | 104.00 |
| C2 | 0.30 | 123 | 19103 | 80.90 |
| C3 | 1.00 | 409 | 16584 | 27.70 |
